# Exploring the Impact of Various Treatments on Gene Expression in Olive (*Olea europaea* L.) Drupes Affected by *Phytophthora oleae*: Insights from RNA sequencing-based transcriptome analysis

**DOI:** 10.1101/2024.07.08.602446

**Authors:** Sebastiano Conti Taguali, Mario Riolo, Federico La Spada, Giuseppe Dionisio, Santa Olga Cacciola

## Abstract

*Phytophthora oleae* is a pathogen recently reported to cause fruit rot on olive orchards in Italy and root rot in a natural wild-olive forest in Spain. RNAseq analysis was conducted to gain insight into the molecular mechanisms that trigger a plant defense response upon the inoculation of drupes with *P. oleae* and the pre-treatment with the antagonistic yeast *Candida oleophila* or with culture filtrates of the antagonistic filamentous fungus *Trichoderma atroviride*. Both treatments were applied to the olive drupe 24 h before the inoculation with the pathogen. Although no full resistance was observed, the virulence of *P. oleae* was reduced when the drupes were co-inoculated with the yeast or treated with culture filtrates of *Trichoderma*. Severity of *Phytophthora* rots in olive drupes was assessed at 24, 72, and 168 hours post pathogen inoculation (hpi) and rated based on an empirical scale. The most effective in reducing the disease severity of *P. oleae* infection on olive fruit was the treatment with *T. atroviride* filtrate (56% reduction), followed by *C. oleophila* (52%). Results showed that 2,466, 1,883, and 1,757 genes were differentially expressed in response to *P. oleae*, to the binary pathosystem *C. oleophila* and *P. oleae*, and *T. atroviride* and *P. oleae*, respectively, as compared to wound. Differential RNAseq by DESeq2, performed at 72 hours post-inoculation, and qPCR validation, at 24, 72, and 168 hpi, of the top differentially expressed genes defined a new pattern of plant defense mechanisms involving both PAMP and ETI immunity, with production of ROS and PRs.

## 1. Introduction

*Phytophthora oleae,* the only known species comprised in subclade 2d of *Phytophthora* clade 2d^1,2^, was for the first time reported as an emergent pathogen of olive (*Olea europaea* L.) in southern Italy^3^. Infection by this oomycete causes a soft rot of mature drupes whose symptoms often overlap with those of anthracnose caused by diverse *Colletotrichum* species, which is regarded as the most serious disease of olive fruit worldwide^4–8^. Other *Phytophthora* species were reported to cause rots of olive fruit^9^. Pre- and post-harvest rots of olive drupes incited by fungi and oomycetes may cause severe yield losses and have detrimental effects on oil quality^10–12^. Management strategies to prevent spoilage of olive production, caused by infectious diseases of fruit, have mainly focused on anthracnose. Although those strategies have prevalently relied on the use of synthetic fungicides, alternative means, such as the natural substances and biological control agents (BCAs) have been also considered^4,13,14^. By contrast, minor fungal diseases and emerging diseases of olive fruit caused by oomycetes have so far received little attention. The search for more sustainable alternatives to conventional synthetic fungicides is a new frontier of modern plant pathology. In the olive oil supply chain, the objective of substituting or reducing the use of synthetic fungicides is of particular relevance due to the risk that residues on fruit pass into the oil during the extraction process. It is noteworthy the olive fruit itself is a reservoir of microorganisms that can be exploited as BCAs^15,16^. Among BCAs, *Trichoderma* species have been extensively tested against pre- and post-harvest fruit pathogens, including diverse species of *Phytophthora*^17–23^. It has been also envisaged the possibility of using bioactive metabolites produced by *Trichoderma* species as natural fungicides^22,24^. In field applications, the efficacy of metabolites would be less conditioned by environmental factors than the living microorganism. A more rapid effect would be an additional advantage of metabolites over the living microorganism, as in the olive oil supply chain the post-harvest phase preceding the oil extraction is very brief. Yeasts are other effective BCAs of post-harvest fruit diseases^25–27^. A very interesting antagonistic yeast is *Candida oleophila*, which has showed a biocontrol activity against several post-harvest diseases of diverse fruit crops. Strains of this yeast have been exploited as commercial bioproducts^19,28–30^.

Several lines of evidence indicate multiple modes of action contribute to the effectiveness of BCAs in suppressing plant diseases, including the elicitation of plant immune system^31^. However, few studies have addressed the transcriptomic analysis of the complex interaction pathogen/host/BCA. *Phytophthora* species are known to secrete a large array of effectors during infection of the plant host. In recent decades, significant progresses have been made in identifying numerous *Phytophthora* effectors, broadly categorized into two main types: CRN and RxLR effectors.^32^ The CRN proteins are a family of effectors that cause necrosis in the cells of the host and also induce further intracellular effectors that target the host nucleus during infection. RxLR effectors play a central role in pathogen–plant interactions. They are secreted proteins that manipulate the host’s immune responses, allowing the pathogen to establish infection^33,34^. Although several studies have already elucidated the role of CRN and RxLR proteins in several *Phytophthora* spp., there is a lack of knowledge about the role of these effectors in the pathosystem *P. oleae*/olive drupes.

Successful *Phytophthora* pathogens have evolved and adapted to a specific host range, developing a set of effector proteins and phytotoxins that either directly induce plant cell death or suppress pattern-triggered immunity (PTI) and trigger susceptibility of the host. Nevertheless, some pathogenesis-related proteins (PR-proteins) are constitutive in some plants and represent a first barrier to the pathogen conveying a sort of innate immunity. Most of the time, even such constitutive levels of PR-proteins might be triggered by the ecto-symbiosis with microorganisms, such as *Trichoderma* spp., or their metabolites known to trigger both growth promotional effects and plant defense mechanisms by the elicitation of salicylic acid (SA)-, ethylene (ET)-, and jasmonic acid (JA)-dependent processes^22,35^.

The aims of this study were: i. to evaluate the effectiveness of either *Trichoderma atroviride* culture filtrates or *C. oleophila* strain O in suppressing rot of olive drupes inoculated with *P. oleae*; ii. to examine the differential gene expression in *P. oleae* and olive drupes, treated/non treated with *C. oleophila* or *T. atroviride*-culture filtrate by high throughput technologies (*i.e.* Illumina RNA sequencing).

## 2. Results

### 2.1. Evaluating the effectiveness of *C. oleophila* strain O and *T. atroviride*-culture filtrate in controlling the rot of olive drupes incited by *P. oleae*

This section reports the results of tests aimed at evaluating the effectiveness of either *C. oleophila* strain O cell suspension or *T. atroviride*-culture filtrate in controlling the rot of olive drupes inoculated with *P. oleae* isolate VK10A. Overall, drupes treated with *T. atroviride*-culture filtrate (IDB) or *C. oleophila* cell suspension (IDC) reported values of disease severity significantly lower than control drupe treated with sterile distilled water (sdw) (IDA) (Figure 1). In particular, the mean value of the relative area under the disease progress curve (rAUDPC) was 0.43 for the control (IDA), 0.21 (i.e. 52% reduction compared to the control) and 0.19 (i.e. 56% reduction compared to the control) for the drupes treated with *C. oleophila* (IDC) and *T. atroviride-*culture filtrate (IDB), respectively. *In vitro* inhibitory activity of liquid culture extracts of *T. atroviride* against *Phytophthora* species (*P. nicotianae* and *P. parvispora*) had been demonstrated in a previous study^22^. By contrast, no previous studies reported any data on the efficacy of *C. oleophila* against *Phytophthora* species.

**Figure 1.**
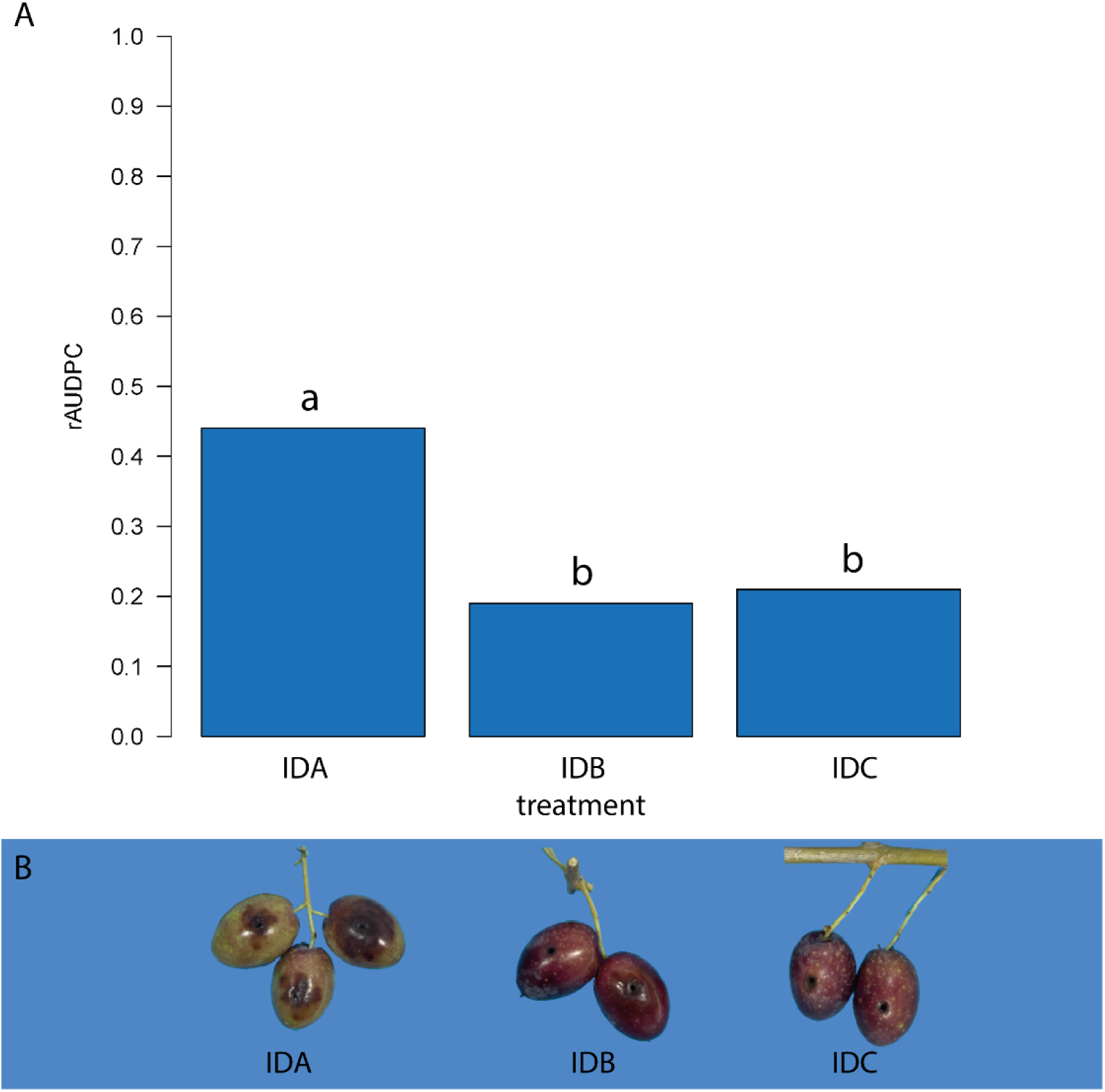
Mean values of the relative area under the disease progress curve (rAUDPC) in olive drupes inoculated with *Phytophthora oleae* and treated with sterile distilled water (control) (IDA), *Candila oleophila* suspension (IDB) or *Trichoderma* atroviride-culture filtrate (IDC). Panel A, rAUDPC values of treatments IDA, IDB and IDC calculated on the basis of % rot severity value recorded at 24, 72 and 168 hours after inoculation (hpi). Values sharing the same letter do not statistically differ according to Tukey’s honestly significant difference (HSD) test (*p* ≤ 0.05). Panel B, Drupes at 72 hpi showing the different size of necrotic lesion: (IDA) wounded drupes inoculated with *P. oleae*; (IDB) wounded drupes inoculated with *P. oleae* and treated with culture filtrate of *T. atroviride*; (IDC) wounded drupes inoculated with *P. oleae* and *C. oleophila*.

### 2.2. Differential RNA analysis at 72 hours post inoculation with *P. oleae*

This section describes the transcriptomic responses of both olive drupes and *P. oleae* in the system olive drupe-*P. oleae*-*C. oleophila* or olive drupe-*P. oleae*-*T. atroviride*-culture filtrate. To this purpose, the following seven treatments were established: (i) non-wounded drupes treated with 20 µl of sdw (ID1); (ii) wounded drupes treated with 20 µl of sdw (ID2); (iii) wounded drupes inoculated with *P. oleae* (ID3); (iv) wounded drupes treated with *T. atroviride-*culture filtrate and inoculated with *P. oleae* (ID4); (v) wounded drupes treated with *C. oleophila* and inoculated with *P. oleae* (ID5); (vi) wounded drupes treated with *T. atroviride*-culture filtrate (ID6); (vii) wounded drupes treated with *C. oleophila* (ID7). Pre-treatment with *C. oleophila* cell suspension or *T. atroviride-*culture filtrate was performed 24h before *P. oleae* inoculation. Drupe fragments were collected at 24-, 72- and 168-hours post inoculation of *P. oleae* (hpi). However, as in the experiment described at paragraph 2.1., 72 hpi was found to be the best timing for the RNAseq evaluation, as the symptoms of *P. oleae* infection were evident (values of rot severity of about 25%).

The following paragraphs describe the RNAseq analysis of olive fruit performed by analyzing binary conditions (see paragraphs: comparisons 1, 2, 3, 4, 6 and 7). A cluster analysis was also carried out among all the treatments for the olive genes in all the conditions analyzed (see paragraph: comparison 5). Finally, a RNAseq analysis was carried out in the same samples to highlight the differential mRNA changes of *P. oleae* versus the different treatments with either *C. oleophila* cell suspension or *T. atroviride-*culture filtrate; to this aim, pairwise comparisons (see paragraphs: comparisons 8 and 9) or PCA plots were elaborated.

#### 2.2.1. Comparison 1: intact olive drupes (treatment ID1) vs. wounded drupes treated with sterile distilled water (treatment ID2)

Only 243 (192 upregulated and 51 downregulated) differentially expressed genes (DEGs) of the olive fruit were detected after filtering with a false discovery rate (FDR) p-value of 0.001 (Figure 2). The upregulated genes included enzymes involved in secondary metabolite production, particularly berberine bridge enzymes (BBE; LOC111391703 and LOC111391702), a subgroup of the superfamily of FAD-linked oxidases that could either be involved in isoquinone alkaloid biosynthesis or in the oxidation of carbohydrates at the anomeric center to the appropriate lactones. Besides, oligogalacturonides (OGs), small fragments coming from the hydrolysis of pectin-homogalacturonan, represent the best example of damage-associated molecular patterns (DAMP) which activate immunity^36^. However, once the DAMP immunity has been established, the excess of OG accumulation has a negative impact on the plant recovery, which could lead to cell death. BBEs are probably involved in oxidizing oligosaccharide fragments that might be released during mechanical wounds or pathogenesis and could act as DAMPs^37^. Thus, BBE can efficiently neutralize the OGs by oxidation to the lactone counterparts^38^, terminating the DAMP activation response^39^.

**Figure 2.**
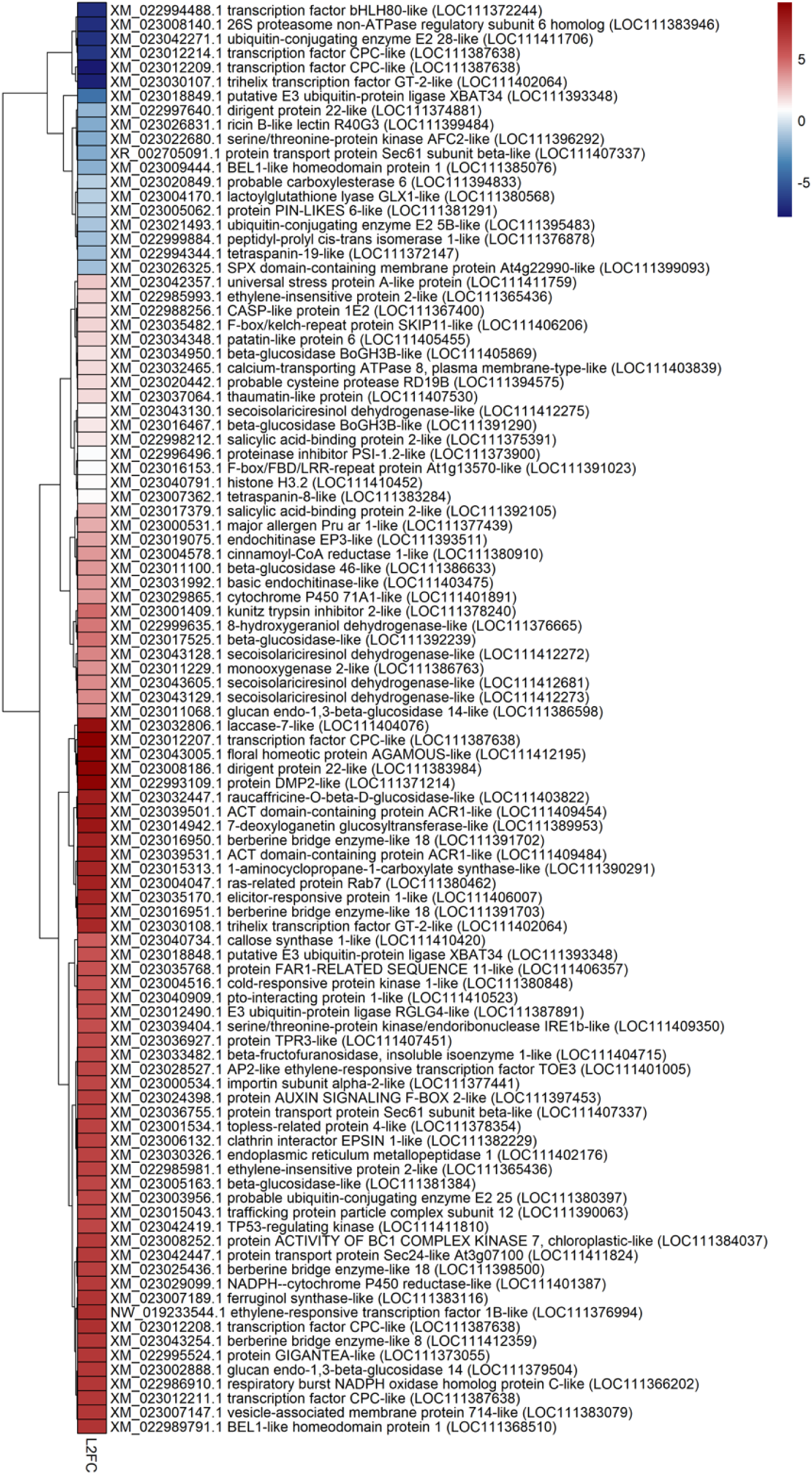
Selection of top up/down-regulated genes present at 72h post mechanical wounding in mature olive cv. Coratina fruits versus intact drupes. The DESeq2 algorithm has been used to calculate the statistics and the differentially expressed genes were sorted according to the log2FoldChange (L2FC). Filtering by FDR with p-value of 0.01 has been applied. Heatmap colors with similar LF2C have been clustered. The complete set of up-down-regulated genes is present in **Supplementary Table 1**.

Other top DEGs include the ethylene biosynthetic isogenes of 1-aminocyclopropane-1-carboxylate synthase-like (ACC synthases; LOC111390291, LOC111392037, and LOC111370655). Indeed, their enzyme activity is involved in the synthesis of the precursor, 1-aminocyclopropane 1-carboxylic acid (ACC), synthesized from S-adenosyl-L-methionine (SAM) by ACC synthases (ACSs) and finally oxidized to ethylene (ET) by ACC oxidases (ACOs)^40^. It is known that gaseous ET can trigger and enhance host resistance and basal resistance to fungal pathogens^41^. The ET perception and activation of the defense mechanism pass through the ET receptors and ET transcription factors (TFs) that mediate the signal transduction until the ethylene-insensitive EIN TF (EIN2 and EIN3) hubs mediated by the F-box genes (e.g. EBF1 and EBF2)^42^. ET receptors are represented by a family of receptors localized in the (ER) membrane. Two main types of ethylene receptors are known; Ethylene response1 (ETR1) and ERS1, belong to subfamily 1c, and ETR2, EIN4, and ERS2 belong to subfamily 2^43^. Other TFs are activated by ET, such as APETALA2-ethylene responsive factors (AP2-ERFs) which are mediators of stress responses and developmental programs^44^. Many ET-related TFs here were DEGs as the ETR TF 1B-like (LOC111378715) which belong to the AP2 DNA-binding TFs in plants such as APETALA2 and EREBPs^45^. The elicitin INF-1 of *P. infestans* induced several defense responses in *Nicotiana* spp., including reactive oxygen species (ROS) and ET production with consequent hypersensitive cell death (HCD) and accumulation of the sesquiterpenoid phytoalexin capsidiol^46^. In this specific case, the production of phytoalexins is regulated by the AP2/ERF TF, NbERF-IX-33^46^.

Dirigent (DIR) protein 22-like (LOC111383984) was also a highly upregulated DEG, probably involved in catalyzing regio- and stereoselectivity of bimolecular phenoxy radical coupling during lignin or lignan biosynthesis. Dirigent proteins in plants modulate cell wall homeostasis during abiotic and biotic stress^47^. Another top DEG, which is typically expressed after wounding, is beta-amyrin 28-oxidase-like (LOC111396592), a cytochrome CYP716A244 which is implicated in the biosynthesis of oleanane-type saponin biosynthesis^48^. Beta-amyrin synthase and beta-amyrin oxidase were characterized in a yeast, where they were able to form not only α-amyrin 6, but also β-amyrin 1, butyrospermol 23, and Ψ-taraxasterol 19^48^. Nonsterol triterpenoids of *O. europaea* are then metabolized into multioxygenated compounds and these could represent the precursors of triterpene saponins^49^. Triterpenoids as well as saponins isolated from various plant organs from different species are known to possess antimicrobial as well as antioxidant activities^50^.

Among top DEGs, it is noteworthy the presence of elicitor-responsive protein 1-like (LOC111406007), which encodes a protein possessing C2 domains which confer a Ca^2+^-dependence or binding activity. Usually, such C2 domains bind a wide variety of substances including bound phospholipids, inositol polyphosphates, and intracellular proteins probably acting as signal transduction enzymes. The excess of calcium release during the wound or infection can disrupt the calcium signaling. In *Arabidopsis*, its homologue, NP_001078296.1, is annotated as a calcium-dependent lipid-binding (CaLB domain) family protein, and it acts as a novel repressor of abiotic stress response by chelating the excess free Ca^2+^ ^51^.

Among the innate immunity activated defense genes/proteins which have been detected here as DEGs are protease inhibitors (e.g., glu, ase, PSI, potato proteinase inhibitor I, kunitz trypsin inhibitor 2), endo-1,3-beta-glucosidases, endo- and exo-chitinases, pathogenesis-related protein STH-2-like, thaumatin-like protein, cysteine proteinase inhibitor B-like, and defensin-like protein (Supplementary Table 1).

#### 2.2.2. Comparison 2: *Phytophthora oleae*-inoculated wounded drupes (treatment ID3) vs. sterile distilled water-treated wounded drupes (treatment ID2)

About 2466 (854 upregulated and 1612 downregulated) genes were differentially expressed with a FDR p-value of 0.001; top upregulated genes are shown in Figure 2. However, when differentially expressed genes are analyzed by the DESeq2 algorithm, a base PFKM mean is also given. Both log2FoldChange and the baseMean of the FPKM counts are not high enough to have a real biological significance in the differential RNAseq data. Therefore, both log2FoldChange (L2FC) and PFKM (baseMean) have been further filtered by a baseMean value above 2.5 (Figure 3).

**Figure 3.**
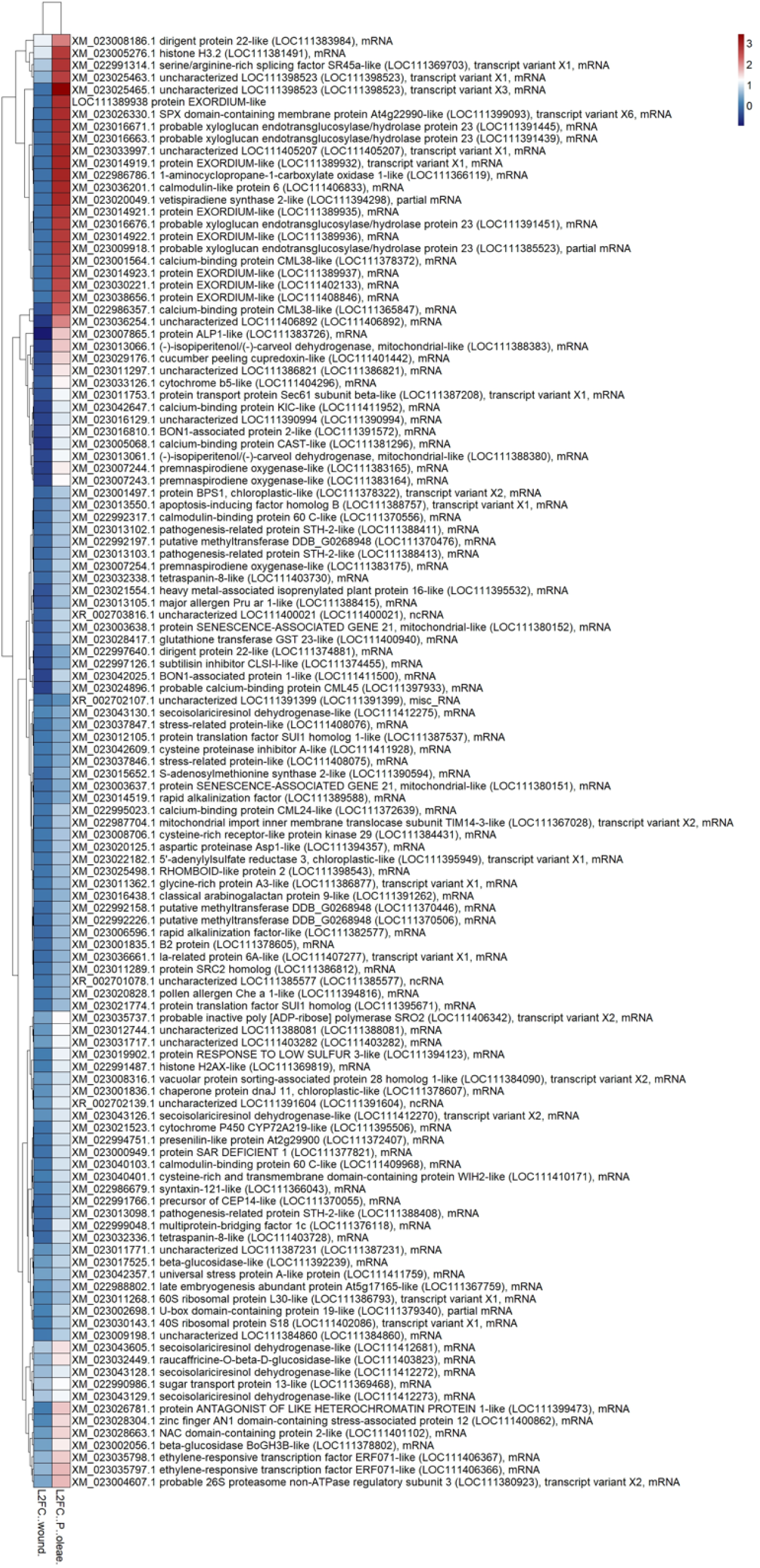
Heatmap of filtered mRNA DEGs present at 72 h post *P. oleae* inoculation of wounded drupes vs. non-inoculated wounded drupes. The wounded L2FC has been calculated between wounded vs not wounded drupes. The DESeq2 algorithm has been used to calculate the statistics and the DEGs were sorted and clustered according to *P. oleae* infection triggering the highest olive mRNAs (positive values L2FC). Filtering by FDR with p-value of 0.001 has been applied. Heatmap shows filtered mRNA as FPKM > 50 as baseMean. The complete set of up- and down-regulated genes is shown in **Supplementary Table 2**.

The top differentially expressed transcripts were mRNA encoding many EXORDIUM-like proteins (e.g. LOC111408846, LOC111402133, LOC111389937, etc.), presumably involved in cell proliferation. EXORDIUM proteins were first identified as mediators of brassinosteroid (BR)-promoted growth in *Arabidopsis*^52^, but they can be negatively regulated by exogenous cytokinin^53^. EXORDIUM 2 like (EXL2, Glyma15g06010) was upregulated and classified among the incompatible interaction genes (IIGs) in the case of *P. sojae* infecting soya bean near isogenic lines (NILs) and associated with resistance^54^.

Another differentially expressed gene was the BON1-associated protein 2 (LOC111391572) which is part of the regulatory complex formed by the BONZAI1 (BON1):BON1-associated protein (BAP1) proteins^55^. BON1-associated proteins seem to exert a negative regulator role related to cell death and defense responses. In plants, BON1 and BAP1 protein-protein interaction sends a negative activation signal to a suppressor of npr1-1, constitutive 1 (SNC1), a nuclear R protein^56^. The product of R gene SNC1 represents a hub of different signaling events coming from many different protein receptor kinases at the plasma membrane, which mediate the final event leading to the activation, or not, of plant PTI or effector-triggered immunity^56^.

Ethylene biosynthesis is also highly upregulated in the case of *P. oleae* infection, since the final enzyme involved in its biosynthesis, the 1-aminocyclopropane-1-carboxylate oxidase 1-like (ACO, LOC111366119), is markedly upregulated as compared to wounded fruit treated with sdw (L2FC≥ 7.6). Two other ACO isogenes were detected as differentially expressed genes (e.g. LOC111370655 LF2C≥4 and LOC111370659 L2FC≥1.9), suggesting that this final step of ET biosynthesis occurs to a high degree during infection^57,58^.

As mentioned before, Ca^2+^ signaling is very active during wounding. Coupled with pathogen attack, Ca^2+^ signaling is further amplified, as inferred by the fact that a higher number of upregulated calcium binding proteins are differentially expressed during *P. oleae* infection: calcium-binding protein CML38-like (LOC111365847), calmodulin-like protein 6 (LOC111406833), calcium-binding protein CML19 (LOC111378369), probable calcium-binding protein CML30 (LOC111406832), calcium-binding protein PBP1-like (LOC111402783), calmodulin-binding protein 60 C-like (LOC111409968), and others. In particular, calmodulin-binding protein 60 C-like (CBP60, LOC111409968) and calmodulin-binding protein 60 C-like (LOC111370556), as well as the transcription activator SAR DEFICIENT 1-like (LOC111402349) and protein SAR DEFICIENT 1 (SARD1) (LOC111377821) are known to bind directly to the promoter of ISOCHORISMATE SYNTHASE 1 (ICS1), increasing the production of salicylic acid (SA) and/or binding to the promoter of ABERRANT GROWTH AND DEATH2 (AGD2)-LIKE DEFENSE RESPONSE PROTEIN 1 (ALD1) and SAR DEFICIENT 4 (SARD4), which leads to the biosynthesis of pipecolic acid (Pip)^59^. In turn, SA and Pip lead to the activation of immune-related genes, such as PATHOGENESIS-RELATED GENES 1 (PRs)^60^. CBP60 is also known to bind the promoter region of SARD1, suggesting that it directly promotes SARD1 expression^61^. Activation of PTI and effector-triggered immunity at local infection sites leads to the development of systemic acquired resistance (SAR) in distal parts of the plant, of which SA and Pip represent the signaling molecules^62^.

Another differentially expressed gene, which encodes the protein SRC2 homolog (LOC111386812), is a calcium binding protein localized in the plasma membrane which possesses a C2 domain. This latter is known to be involved in protein–protein interactions, binding of phospholipids, and targeting of proteins to the membrane in response to Ca^2+^ signaling^63^. However, in the case of pepper infection, when *Phytophthora capsici* secretes a protein elicitor 1 (PcINF1) with homology to the elicitor of the hypersensitive reaction (HR), it induces a cell death response in pepper leaves mediated by the SRC2 protein^64^. In this case, the upregulation of olive SRC2 could represent a negative factor for plant immunity, but act as a positive mediator of the *P. oleae* INF1 effector by increasing its cell death efficacy.

A huge transcriptional remodeling during *P. oleae* infection is suggested by the massive upregulation of olive histones (Supplementary Table 2): histone H3.2 (LOC111381491), histone H2AX-like (LOC111369819), histone H4 (LOC111408016), histone H2AX-like (LOC111379587), and histone H2B.5-like (LOC111399700). Genome remodeling via epigenetics (changes in methylation level) and transposases activity are known to occur during pathogen attack and the development of plant immunity^65^. In plants, acetylation of histones H3 and H4 (H3ac and H4ac), trimethylation on lysine residues, and mono-ubiquitination of H2B (H2Bub1) are normally signs of heterochromatin formation with enhanced transcriptional activation^66^. Transcriptional remodeling and genome rearrangement including retrotransposon jumping within the chromatin play a central role in plant defense mechanisms^67,68^. Based on *de-novo* synthesis of histones alone, it cannot be possible to discern their epigenetic post-translational modification, but certainly, artificial treatment of plants with SA induces acetylation of histones H3 and H4^69^.

Other differentially expressed genes are the product of the genes encoding olive E3 ubiquitin-protein ligases. Some of those could be the homologue of Arabidopsis RING E3 ligase HUB1 which mediates histone mono-ubiquitination with the subsequent activation of heterochromatin and probably contributes to enhancement of the defense mechanism^70^.

That genome rearrangement could be linked to the induction of plant resistance that is evident when DDE-type transposases (containing a DD[E/D]-motif) are involved^71^. In this study, it is highly probable that *P. oleae* induced a high expression of the protein ALP1-like (LOC111383726), which possesses a N-terminal RNase H domain, a DNA binding domain and a known transposase DDE C-terminal domain. Demethylation of transposable elements (TEs) makes them jump in heterochromatin genes, which, in this case, could be associated with mediating a ‘long-term memory’, since their insertion into the promoter of these genes makes them constitutively expressed. This epigenetic stress memory can be transmitted to subsequent generations and involves changes in the silencing of transposable elements (TEs) by DNA methylation, histone modifications, and non-coding RNAs. At late infection stages, the DNA jumping activities of functional class I (‘copy and paste’) and class II (‘cut and paste’) TEs, might result in small and large mutations at the sites of excision (class 2) and insertion (class 1 and 2). TE-induced insertion/excision accelerates the evolution of novel defense regulatory genes and, because TEs are somewhat regulated by epigenetic mechanisms, the derived defense genes will remain under stress-dependent epigenetic control and potentially able to resist to biotic stresses^72^.

Transcription factors (TFs) known to be involved during both wound response and pathogen attack are the WRKY TFs^73^. These TFs contain a 60-70 amino acid protein domain composed of a conserved WRKYGQK motif and a zinc-finger region (C2HC). Both domains contribute to the capability of DNA to bind with W-box and/or other cis-acting elements, such as in R-genes^74^ or biosynthetic genes related to specialized metabolism^75^.

Upon *P. oleae* infection, it has been recorded an upregulated (L2FC≥ 3.4) olive WRKY TF 33 (LOC111405948) homologue of AtWRKY33, which functions as a positive regulator of resistance toward some necrotrophic fungi^76^. However, the olive WRKY TF 18-like (LOC111377241) was even more differentially expressed (L2FC≥ 4.6) and, according to the homology function with AtWRKY18, could enhance certain defense signaling, boosting the transcription of PR genes and resistance against certain pathogens^77^. Other upregulated WRKY TFs include WRKY TF 53 (LOC111367778. L2FC>6.1), WRKY transcription factor 23 (LOC111388449, L2FC>4.7), WRKY transcription factor 70 (LOC111392938, L2FC>4.3), and WRKY transcription factor 57 (LOC111373803, L2FC>1.2).

The main differentially upregulated R-genes were the LEAF RUST 10 DISEASE-RESISTANCE LOCUS RECEPTOR-LIKE PROTEIN KINASE-like 1.1 (LOC111390435, L2FC>5.2), the protein ENHANCED DISEASE RESISTANCE 2 (LOC111389201, L2FC>5), the disease resistance protein At5g63020 (LOC111397442, L2FC>5), receptor-like protein kinase HERK 1 (LOC111407351, L2FC>5.5; LOC111407714, L2FC>2.3), the rust resistance kinase Lr10-like (LOC111411655, L2FC>3.6), the disease resistance RPP13-like protein 1 (LOC111400669, L2FC>3.6), the disease resistance protein RGA1 (LOC111392054, L2FC>3.2), and the disease resistance RPP13-like protein 4 (LOC111379833, L2FC>2.2). These plasma membrane-located receptors possess a nucleotide-binding site–leucine-rich repeat (NBS-LRR) and a protein kinase C domain for intracellular signal transduction and are probably involved in the signaling cascade of pathogen sensing^78^. On the other hand, upregulated F-box/LRR-repeat proteins might be associated with resistance to *Phytophthora* species, since for instance, in cacao (*Theobroma cacao*), similar genes were differentially upregulated in response to *P. megakarya* only in a resistant cultivar^79^. Here, upregulated F-box/LRR-repeat proteins were represented by F-box/FBD/LRR-repeat protein At1g13570-like (LOC111386658) and the F-box/kelch-repeat protein At1g67480-like (LOC111406244) (Supplementary Table 2). Other differentially expressed TFs during *P. oleae* infection are the ethylene-responsive TFs ERF104-like (LOC111386758), ERF017-like (LOC111373863), ERF017-like (LOC111373875), ERF118-like (LOC111403203), ERF071-like (LOC111406367), RAP2-3-like (LOC111403560), ERF4-like (LOC111395465), and ERF04 (LOC111381486). Ethylene induced immunity involves sensing and TF-responsive transcription; this ET-pathway interacts both positively and negatively with the SA-pathway, in relation to the type of plant and pathogen^80^. The APETALA2 (AP2)/ETHYLENE RESPONSE FACTOR (ERF) transcription factor ORA59 is a major positive switch of the ET/JA-mediated defense pathway in *A. thaliana*. In *Arabidopsis*, biosynthetic genes responsible for the formation of hydroxycinnamic acid amides (HCAAs), known as phytoalexins, are regulated by ORA59 and other ET-responsive TFs^81^.

Jasmonic acid (JA) and its precursors and derivatives, referred to as jasmonates (Jas), are important molecules in the regulation of plant responses to biotic and abiotic stresses^82^. JA-dependent defense responses are predominantly effective against necrotrophic pathogens and their downstream signaling pathway is typically targeted by fungal effectors to prevent the expression of JA-related defense genes. Coronatine-insensitive protein 1 (COI1) is an F-box protein essential for all the jasmonate responses. COI1 interacts with multiple proteins to form the SCF:COI1:E3 ubiquitin ligase complex which recruits jasmonate ZIM-domain (JAZ) proteins for degradation by the 26S proteasome. *Phytophthora sojae* uses an RXLR effector, Avh94, to manipulate host JA signaling to promote infection hijacking-JA signaling^83^. Zhao et al. have demonstrated that Avh94 interacts with soybean JAZ1 or JAZ2, which is a repressor of jasmonic acid (JA) signaling^83^. Methyl jasmonate (MeJA)-triggered JAZ1 protein degradation could be inhibited by the proteasome inhibitor MG132, suggesting that JAZ1 is consistently degraded by the 26S proteasome. Avh94 bind JAZ1 and protect it against methyl jasmonate-induced degradation. In turn, the JAZ proteins act as repressors of JA TFs such as AtMYC2, which is involved in the induction of JA biosynthetic genes. In the case of *P. oleae* infection, the coronatine-insensitive protein 1-like (LOC111390662) is a differentially expressed gene and could represent a jasmonate receptor. The highly differential expression of COI1 is accompanied by other upregulated differentially expressed genes encoding F-box proteins: F-box protein At3g07870-like (LOC111409106), F-box protein At1g61340-like (LOC111407155), and F-box protein PP2-A12-like (LOC111408420).

Among differentially expressed genes related to the PR proteins, in the case of *P. oleae* infection, two genes are predominating, the PR proteins STH-2-like (LOC111388408, L2FC≥5) and STH-1-like (LOC111388885, L2FC≥2.3). LOC111388408 was a gene upregulated in response to wounding, but it is even more upregulated in response to the pathogen. Furthermore, another upregulated gene, as compared to the wound condition itself, is the cysteine proteinase inhibitor B-like (LOC111390855, L2FC≥6) and the cysteine proteinase inhibitor A-like (LOC111411928, L2FC>3) which could act in defense, trying to block several *P. oleae* cysteine proteases. An upregulated gene related to the apoplastic immunity/defense protease was the cysteine protease RD19B (LOC111394575) of the *O. europaea* homologue of RD19, an *Arabidopsis* cysteine protease that modulates RESISTANT TO RALSTONIA SOLANACEARUM 1-R (RRS1-R)^84^. Other differentially expressed genes related to defense response, from highest to lowest L2FC, respectively, are pollen allergen Che a 1-like (LOC111394816, L2FC≥4.7), subtilisin inhibitor CLSI-I-like (LOC111374455, L2FC≥4.5), and kirola-like proteins (LOC111411103, and LOC111410001). On the other hand, a defense response can be represented by the upregulated expression of proteases such as aspartic proteinase Asp1-like (LOC111394357, L2FC>2.2), subtilisin-like protease SBT1.6 (LOC111374885, L2FC>2.1), and SBT3.15 (LOC111386418, L2FC>1.7).

For most of the common differentially expressed genes detected in this comparison, a higher level of upregulation was generally observed. Importantly, tetraspanin-2 (LOC111399506) and tetraspanin-8-like (LOC111403728, and LOC111383284) are genes related to the wounding condition itself (Figure 1). Tetraspanins are transmembrane specific proteins of extracellular vesicles (EVs) or exosomes (EXOs)^85^, which possess the function of cargo vesicle shipping molecules involved in plant–pathogen interaction. Exosomes transport lipids, proteins (e.g. PR proteins), messenger RNA, and especially microRNAs^86^. Silencing of the pathogens by miRNA is a recent trend in pathogen resistance^87^. Indeed, plant EVs inhibit fungal growth and virulence by transferring their cargo content into fungal cells at the appressoria/haustoria^88^. Another role of tetraspanins is in moving the EVs through infection hot spots and allowing the exosomal NADPH oxidase (NOX) to release ROS molecules locally (e. g. H2O2 production).

It is also worth mentioning the differential expression of the gene encoding for the receptor-like protein kinase FERONIA (FER; LOC111374777, L2FC≥5.5) (Supplementary Table 2). FER negatively regulates plant immunity by inhibiting JA and coronatine (COR) signaling. It phosphorylates and destabilizes MYC2, the master regulator of JA signaling, which otherwise activates SA biosynthesis^89^. FER, in response to abiotic and biotic stress, acts as a sensor of cell wall integrity which could be challenged by pathogen secreted enzymes^90^. FER may negatively regulate plant immunity to biotrophic pathogens^91^. The plant rapid alkalinization factor (RALF) are secreted peptides that were first identified through their ability to trigger a rapid increase in extracellular pH [a]. Their mechanism of action is through binding to FERONIA, that once activated triggers extracellular alkalinization, to favor pathogenic subtiliases (alkaline proteases) or alkaline cysteine proteases and inhibits the defense response^91^.

#### 2.2.3. Comparison 3: wounded olive drupes pre-treated with *C. oleophila* and inoculated with *P. oleae* (treatment ID5) vs. wounded olive drupes inoculated with *P. oleae* (treatment ID3)

A total of 1883 (1018 upregulated and 865 downregulated) olive differentially expressed genes were detected in this comparison (Supplementary Table 3). It is known that *C. oleophila* secretes various cell wall-degrading enzymes, exo-β-1,3-glucanases, chitinases, and proteases, which are especially active against filamentous fungi^92^. *C. oleophila* has an *in vitro* effect on the inhibition of mycelial growth of several *Phytophthora* species^93^. Due to this beneficial effect, some of the genes in *O. europaea* related to the pathogen response could be downregulated, since the pre-treatment ameliorates the pathogenic index (rAUDPC = 0.21; Figure 1, IDC). Thus, the downregulated genes, in this comparison, could represent the olive genes that help to counteract the pathogen invasion, or which are no longer needed since the pathogenic effect of *P. oleae* decreased due to *C. oleophila* pre-treatment. Among the top downregulated genes with L2FC>-21 were ALA-interacting subunit 1-like (LOC111368552), the ras-related protein Rab7 (LOC111380462), and protein YIF1B-like (LOC111391136), all localized into the Golgi apparatus, regulating endomembrane dynamics and involved in secretory vesicle formation^94^. In the specific case here, RABs, which are the largest family of small guanosine triphosphate (GTP)-binding proteins, are involved in intracellular trafficking and in autophagy or plant microbe interactions, and in biotic and abiotic stress responses^94^, for instance, the secretion of PATHOGENESIS-RELATED 1 (PR1) protein to the apoplast. *Phytophthora capsici* RxLR effector RxLR242 has been found to bind RABE1-7, a ras-related protein, and inhibits vesicle trafficking^95^. This detected decrease of vesicle trafficking is accompanied by the strong downregulation of the WRKY TF 18-like (LOC111377241) here with L2FC>-8.5 instead of +3.5 as for *P. oleae* infection itself. Otherwise, WRKY 18, as mentioned, could have further enhanced another defense signaling^75^.

One of the possible mechanisms by which *O. europaea* counteracts *P. oleae* infection is through synthesis of sesquiterpenoid phytoalexins. In potato challenged by *P. infestans*, it has been found that the infected tissue produces antimicrobial sesquiterpenoid phytoalexins (*e.g.* lubimin and rishitin) which might have been implicated in a resistance mechanism in some cultivars^96^. Here, there is a clear downregulation of sesquiterpenoid phytoalexin biosynthesis when drupes are pre-treated with *C. oleophila* and inoculated with *P. oleae* (treatment ID5) as compared to drupes inoculated with *P. oleae* (treatment ID3) (Supplementary Table 3). Vetispiradiene synthases or sesquiterpene cyclases and cytochrome P450 premnaspirodiene oxygenases catalyze several key biosynthetic steps in the synthesis of antimicrobial (phytoalexins) sesquiterpenes^97^. Indeed, the expression of olive vetispiradiene synthase 2-like (LOC111371938, L2FC>-7.1), vetispiradiene synthase 2-like (LOC111372151, L2FC>-7), premnaspirodiene oxygenase-like (LOC111369993, L2FC>-6.5; LOC111383354, L2FC>-5.9; LOC111375139, L2FC>-5.2; LOC111374442, L2FC>-4.6), vetispiradiene synthase 2-like (LOC111394812, L2FC>-6.3), vetispiradiene synthase 2-like (LOC111394298, L2FC>-5.7), and (-)-isopiperitenol/(-)-carveol dehydrogenase (LOC111388383, L2FC>-5.7; LOC111388380, L2FC>-5.0) clearly points to the production of sesquiterpenoid defense compounds. These enzymes are upregulated in treatment with the only inoculation of *P. oleae* (ID3) (Supplementary Table 3) but also during *P. infestants* infection in potato. Associated with decreased infectivity of *P. oleae* due to *C. oleophila* pre-treatment, production of these phytoalexins is downregulated. However, the amount of phytoalexins produced also depends on the amount of the precursor mevalonate and hence, the expression level of 3-hydroxy-3-methylglutaryl CoA reductase (HMGR)^98^.

Lignan biosynthesis is also regulated differentially from that occurring during wounding and plant defense responses in this comparison. Many phytoalexins in plants are represented by complex dimeric substituted lignan^99^. Dimerization of two coniferyl alcohol units produces pinoresinol, which is then reduced to lariciresinol and secoisolariciresinol by pinoresinol/lariciresinol reductase (PLR), and finally into matairesinol by secoisolariciresinol dehydrogenase (SIRD)^100^. Here, opposite to what was observed in olive drupes only inoculated with *P. oleae* (treatment ID3) (Supplementary Table 2), the biosynthetic enzymes of the lignan biosynthesis are also downregulated with *C. oleophila* co-infection: dirigent protein 22-like (LOC111374881, L2FC>-3.5), dirigent protein 19-like (LOC111376314, L2FC>-3), dirigent protein 22-like (LOC111383984, L2FC>-2), and the secoisolariciresinol dehydrogenase-like (LOC111412270, L2FC>-3.6; LOC111412278, L2FC>-1.3) (Supplementary Table 3).

On the other hand, *C. oleophila* pre-treatment also triggered the upregulation of beneficial genes/proteins that help to counteract the *P. oleae* infection. For instance, the PR proteins STH-21-like (LOC111388400) and STH-2-like (LOC111388410) showed L2FC≥1.09 and =1.7, respectively, but baseMean (PFKM) counts of 4032 and 2646. This means that even a small log2-fold increase with elevated high expression counts could be significant if the *p-*value and *p-*adj value are small. The lowest *p-*value and *p*-adj value of the upregulated mRNA/genes during *C. oleophila* co-infection were also filtered by high baseMean and L2FC (Figure 4). Many differentially expressed genes related to defense proteins are also upregulated, such as proteinase inhibitor PSI-1.2-like (LOC111373900), xylanase inhibitor thaumatin-like protein 1b (LOC111367584), chitinase hevamine-A-like (LOC111402962), basic endochitinase-like (LOC111379188), aquaporin PIP1-3-like (LOC111408442), TIP1-1 (LOC111403122), and PIP2-2-like (LOC111372815) (Figure 4, Supplementary Table 3). The involvement of the aquaporins (AQPs) is not immediate, but it has been shown that they also transport H2O ^101^. Hydrogen peroxide acts as a ROS signaling molecule and can be produced by various oxidases, superoxide and superoxide dismutase, plasma membrane NADPH oxidases, and peroxisomal oxidases, by electron transport in chloroplasts and mitochondria, or by other apoplastic oxidases. In wheat (*Triticum aestivum* L.), TaPIP2;10 transports the pathogen-induced apoplastic H2O2 into the cytoplasm of the infected cell and this signal induces PTI immunity^102^. Consequently, in this comparison, the olive AQPs expressed here might have a function of stimulating the H2O2 signaling and accelerate the PTI response. Besides, the non-symbiotic hemoglobins have been implicated in the scavenging of nitric oxide signaling^103^. In this comparison, the possible role of the upregulation of the non-symbiotic hemoglobin 2-like (LOC111376313) could be that of neutralizing the excess of NO produced during the pre-treatment (Supplementary Table 3).

**Figure 4.**
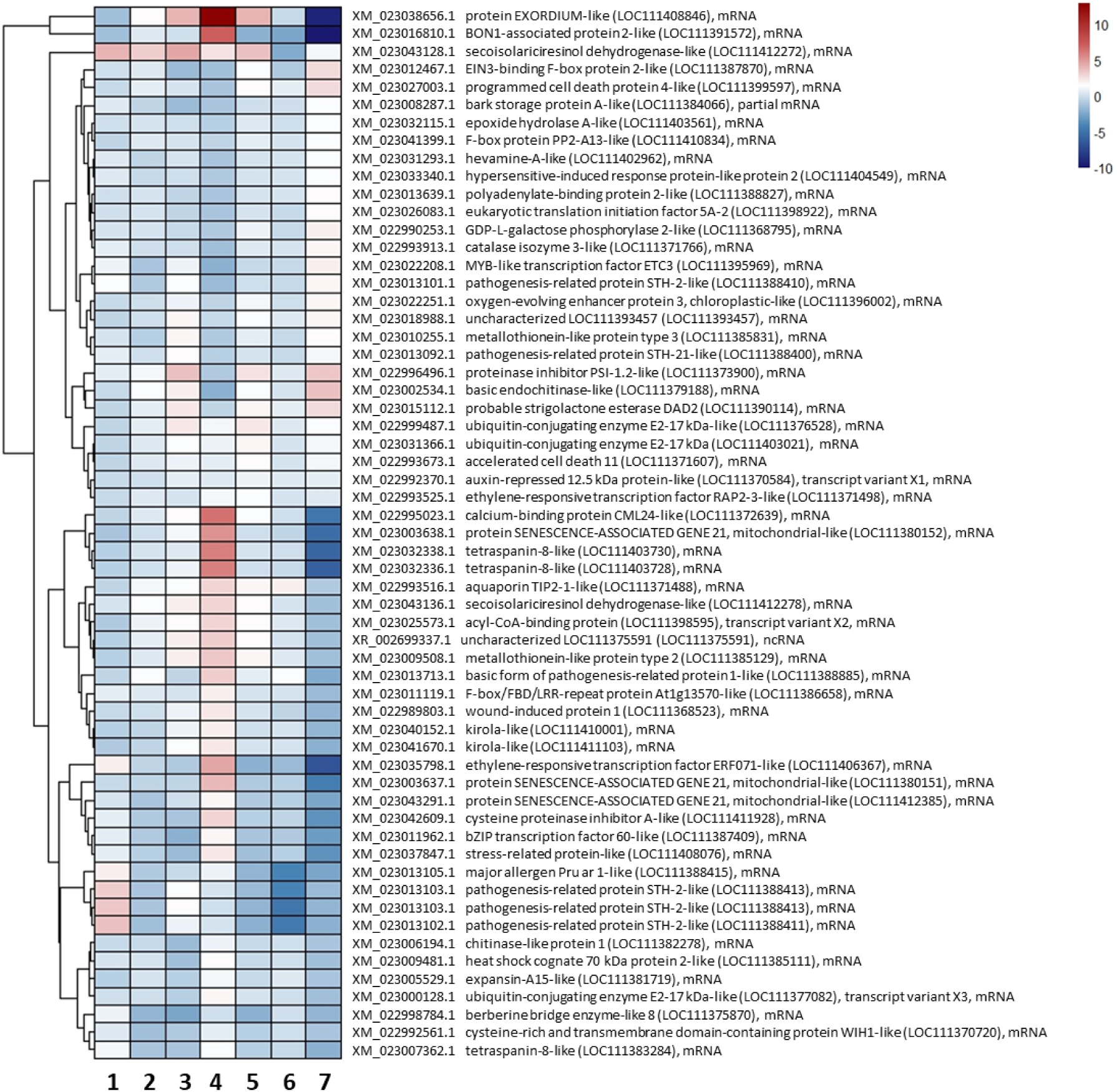
Cluster heatmap analysis of the top mRNAs related to DEGs of *O. europaea* detected 72 hpi from the significative upregulated genes during *C. oleophila* co-infection with *P. oleae* vs. infection with *P. oleae* alone (lane 7). Filtering after DESeq2 analysis was applied to the DEGs in Supplementary Table 3 by sorting for the lowest *p-*value/*p*-adj-value and highest baseMean or L2FC. Lane 1, L2FC of wounded drupes vs. intact drupes; lane 2, L2FC of wounded drupes with *T. atroviride* extract vs. wounded drupes; lane 3, L2FC of wounded drupes plus *C. oleophila* vs. wounded drupes; lane 4, L2FC of wounded drupes plus *P. oleae* vs. wounded drupes; lane 5, L2FC of wounded drupes plus *C. oleophila* and *P. oleae*; lane 6, L2FC of wounded drupes plus *T. atroviride* extract plus *P. oleae* vs. wounded drupes; lane 7, L2FC of wounded drupes plus *C. oleophila* and *P. oleae* vs. wounded drupes and *P. oleae*.

#### 2.2.4. Comparison 4: wounded olive drupes pre-treated with *Trichoderma atroviride*-culture filtrate and inoculated with *Phytophthora oleae* (treatment ID4) versus wounded drupes inoculated with *P. oleae* (treatment ID3).

A total of 1757 (914 upregulated and 843 downregulated) olive DEGs differentially expressed genes were detected in comparison ‘olive drupes pre- treated with *Trichoderma atroviride*-culture filtrate (treatment ID4) and inoculated with *Phytophthora oleae*’ versus ‘olive drupes inoculated with *P. oleae* (treatment ID3)’ (Supplementary Table 4). Biocontrol fungi, such as *Trichoderma* spp., are known to induce systemic resistance (ISR) and prime their host plants to become more resistant to future attack from pathogenic microorganisms^104^. For example, *Trichoderma viride* produces the peptide antibiotic alamethicin^105^, that forms PM-pores and alters the membrane potential and ion permeability, and acts against other fungi or bacteria^106^. The *T. atroviride*-culture filtrate used in this study was checked by proteomics for evaluating its composition in proteins. To this aim, the culture filtrate was subjected to Liquid Chromatographic – Mass Spectrometry (LC-MS) proteomic analysis and the top abundant proteins were identified (Table 1).

**Table 1.**
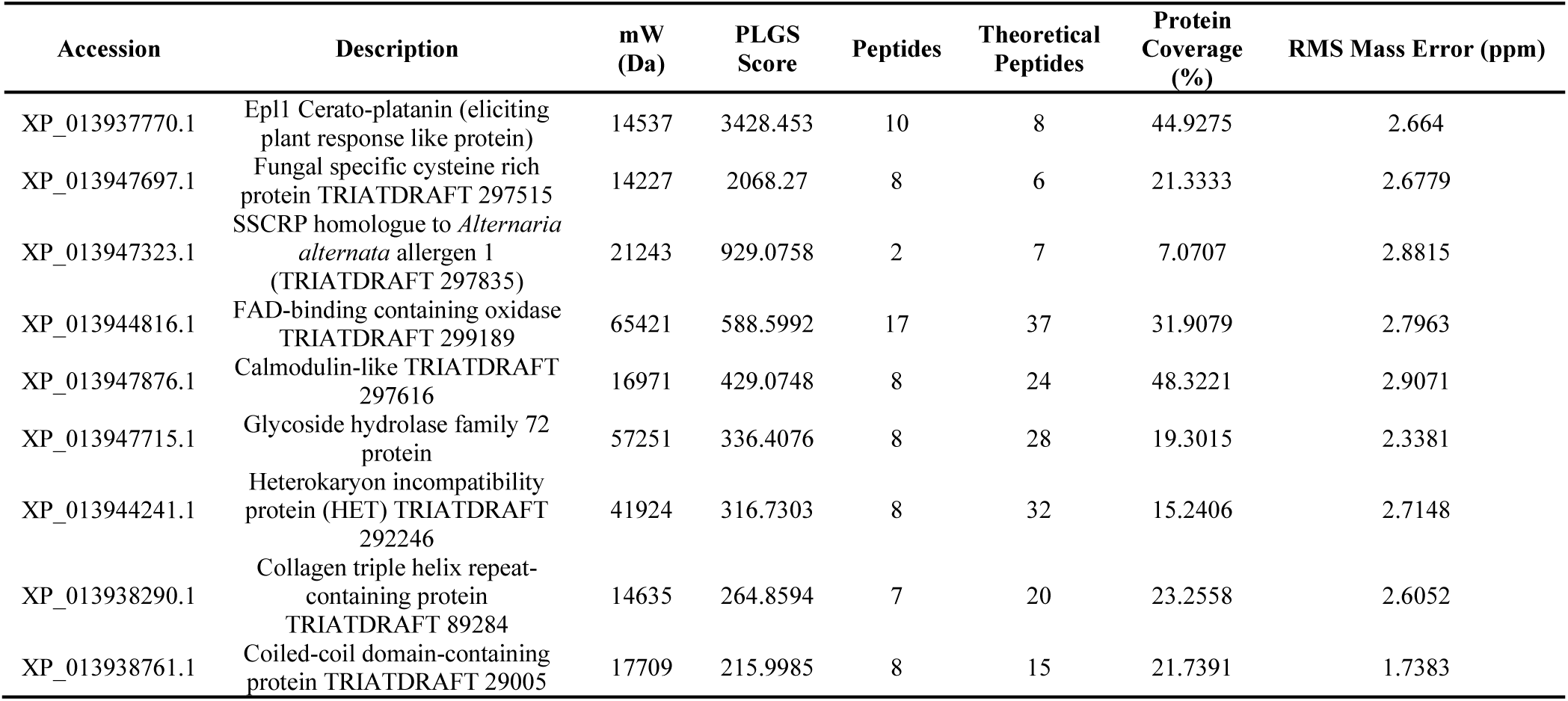
Proteins with the highest concentration in Trichoderma atroviride strain TS-culture filtrate. Identification of the tryptic peptides was made by LC-MS and runs were analyzed by the software Protein Lynx Global Server (PLGS v3.0, Waters, Milford, USA).

The most abundant protein was the Epl1 elicitin, which is a member of the Cerato-platanin family that contains several phytotoxic proteins of about 150 AA residues. Cerato-platanin contains a cysteine-rich domain (four cysteine residues that form two disulfide bonds). *Trichoderma viride* and *T. atroviride* secrete the proteins Sm1 (XP_013943617.1) and Epl1 (XP_013937770.1), respectively, which elicit local and systemic disease resistance in plants^107^. Deletion of epl1 in *T. atroviride* resulted in reduced systemic protection against *Alternaria solani* and *Botritis cinerea*, whereas the *T. virens* sm1 KO strain was less effective in protecting tomato against *Pseudomonas syringae* and *B. cinerea*. Upregulation of epl1 and sm1 led to enhanced basal resistance in tomato i.e., systemic acquired resistance (SAR) and induced systemic resistance (ISR)^108^.

Results from this study suggest that the impact of *T. atroviride*-culture filtrate pre-treatment on *P. oleae* was significative, as it has been observed the lowest reduction in pathogenic index (rAUDPC = 0.19; Figure 1, IDB). Furthermore, RNAseq results strongly suggest that the pre-treatment of olive drupes with this culture filtrate determined a marked reduction of expression of the *P. oleae* infection marker proteins.

The data in Figure 5 many pairwise comparisons have been selected and filtered according to the most significant upregulated differentially expressed genes (lane 7 describe the comparison 4).

**Figure 5.**
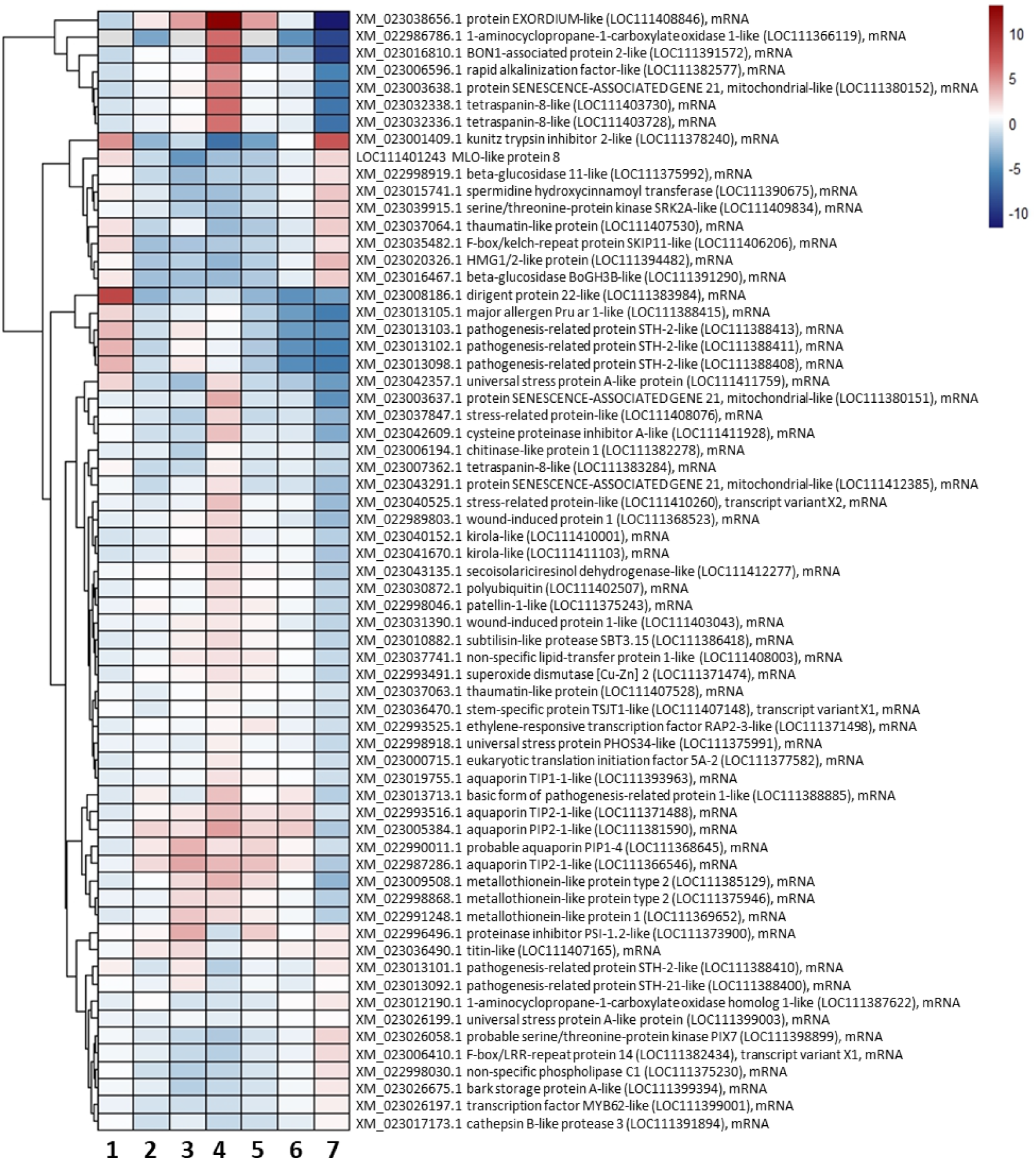
Cluster heatmap analysis of the top mRNAs related to DEGs genes of *O. europaea* detected during 72 hpi from the significant upregulated genes during *T. atroviride culture filtrate* pre-treatment plus *P. oleae* vs infection with *P. oleae* alone (lane 7). The DESeq2 data have been filtered first by lowest pvalue/padj value and then highest baseMean and L2FC from Supplementary Table 4. Lane 1: L2FC of wounded olives vs intact ones; lane 2: L2FC of wounded plus *T. atroviride* extract vs wounded; lane 3: L2FC of wounded plus *C. oleophila* vs wounded; lane 4: L2FC of wounded plus *P. oleae* vs wounded; lane 5: L2FC of wounded plus *C. oleophila* and *P. oleae*; lane 6: L2FC of wounded plus *T. atroviride extract* plus *P. oleae* vs wounded; lane 7: L2FC of wounded plus *T. atroviride* culture filtrate pre-inoculation plus *P. oleae* vs infection with *P. oleae* alone.

By associating differentially expressed genes resulting from comparison 4 with results from LC-MS analysis, it can be speculated that the pre-treatment of olive drupes with of *T. atroviride*-culture filtrate alerts the plant tissues to pathogen attack. Indeed, one of the top upregulated gene, the HR protein-like protein 2, OeHIR-2 (LOC111404549) (Supplementary Table 4), is responsible for alerting the tissues^107^. In this study, two F-box/LRR-repeat proteins 14 (LOC111382434) and (LOC111405202) were also upregulated, but not in stoichiometric amounts as for OeHIR-2 (LOC111404549) (Supplementary Table 4). Therefore, OeHIR-2 should instead be responsible for the disease resistance response, perhaps triggered by *T. atroviride* culture filtrate, which gave the lowest pathogenicity index in this comparison.

Unexpectedly, there were two possibly slightly upregulated resistance genes: LRR, leucine-rich repeat receptor-like serine/threonine-protein kinase homologue of At1g06840 (LOC111371641) and MLO-like protein 8 (LOC111401243) (Supplementary Table 4). Using the Arabidopsis eFP Browser^109^, it is possible to visualize that the expression of At1g06840 mRNA is very high 48 hpi after *Botrytis cinerea* inoculation of leaves. MLO-like protein 8 (LOC111401243) is also overexpressed (Figure 5; Supplementary Table 4).In the barley, the *Mlo* gene, is involved in particular mutation(s) that can cause the misfunction of the MLO protein, resulting in a correlation with resistance to the fungal pathogen *Erysiphe graminis* f. sp. *hordei*^110^. It can be inferred that the same mutations are also present in olive *MLO* germplasm and if those contribute to the partial/total resistance to Oomycetes.

#### 2.2.5. Comparison 5: overall differential mRNA expression of olive genes

Following the pairwise comparisons of olive mRNA in the previous conditions (Figures 2, 3, 4 and 5), it was performed, in addition, the overall changes in mRNA, better summarized by cluster analysis (Figure 6). The following three groups of treatments were used for this cluster analysis: (i) the control groups, including wounded drupes treated with 20 µl of sterile distilled water (treatment ID2), wounded drupes treated with either *T. atroviride*-culture filtrate (treatment ID6) or *C. oleophila* cell suspension (treatment ID7); (ii) olive drupes inoculated with *P. oleae* (treatment ID3); and (iii) *P. oleae*-inoculated olive drupes pre-treated with either *T. atroviride*-culture filtrate (treatment ID4) or *C. oleophila* cell suspension (treatment ID5). In olive drupes inoculated with *P. oleae* (ID3), 739 olive genes were exclusively upregulated (in red in Figure 6) (L2FC > 1) (Figure 6B; Supplementary Table 5).

**Figure 6.**
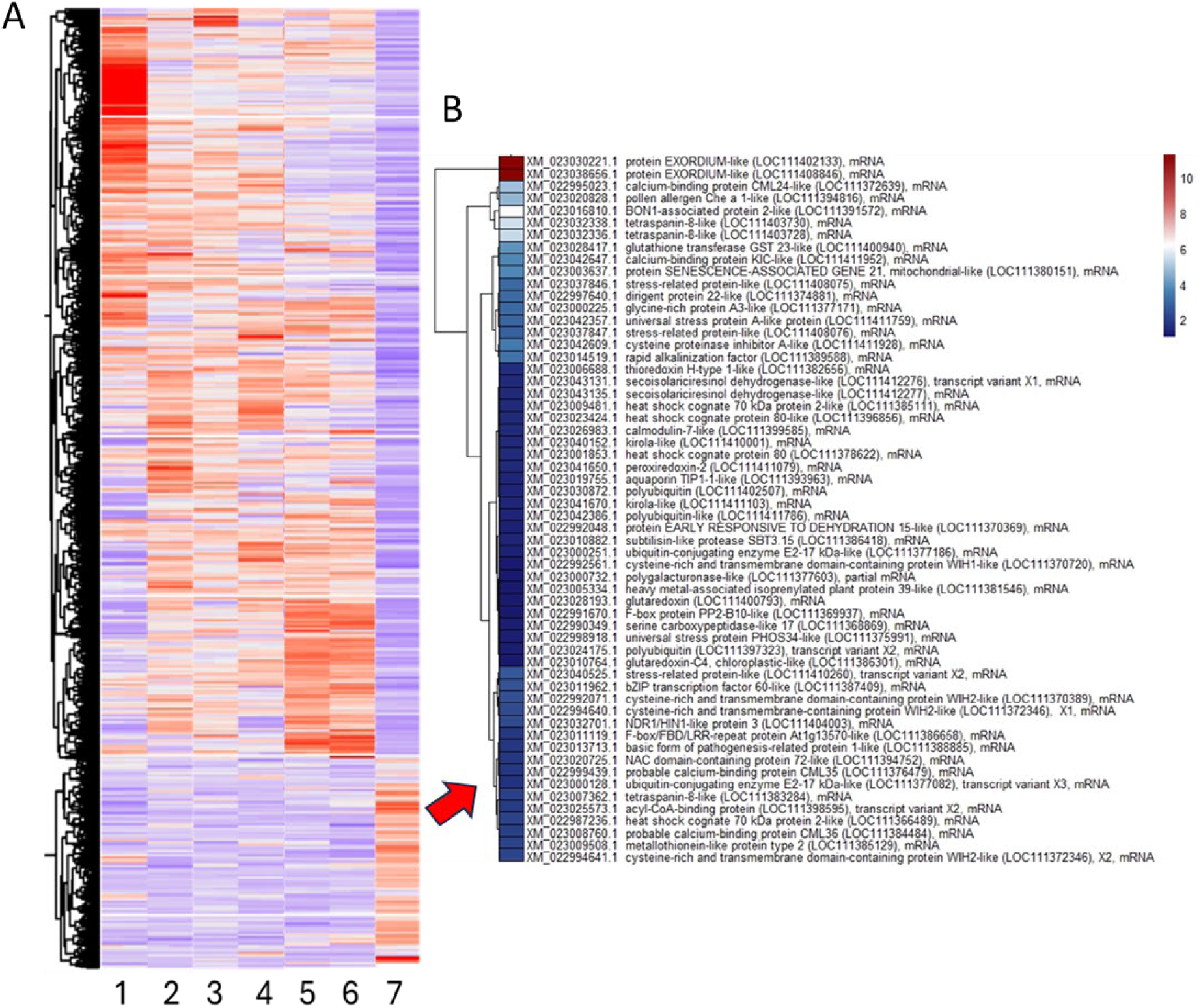
Cluster heatmap analyses of the differential RNAseq related to olive genes among all treatments at 72 hpi. This heatmap has been constructed assuming three groups of samples (all with technical duplicates and some of them with more biological repeats). Panel A (left) cluster analysis in which the wounded drupe treatment LF2C has been normalized to “not wound” ones, while the rest of conditions has been normalized to the “wound” condition. The data has been normalized with L2FC being = +/-4 by Z score. Legend: 1) intact olive drupes (ID1); 2) wounded drupes (ID2); 3) wounded drupes pre-treated with *C. oleophila* cells (ID7); 4) wounded drupes pre-treated with *T. atroviride* culture filtrate (ID6); 5) wounded drupes pre-treated with *C. oleophila* cells and inoculated with *P. oleae* (ID5); 6) wounded drupes pre-treated with *T. atroviride* culture filtrate and inoculated with *P. oleae* (ID4); 7) wounded drupes inoculated with *P. oleae* (ID3). Panel B (right) list of FPKM base mean filtered (>100 counts) upregulated and L2Fc sorted genes in the *P. oleae* alone condition (the total 739 genes are listed in Supplementary Table 5).

The general trend of top upregulated genes triggered by *P. oleae* are described above (Figs. 3 and 4). However, the plant response to the attack of the oomycete, even though this strain of *P. oleae* could not be fully blocked, showed a differential defense response as indicated by the relative expressions of the pathogenic related (PR) proteins (Supplementary Figure 1).

The expression level of PR proteins has been evaluated based on their L2FC (Supplementary Table 6), and their overall comparison are shown in Supplementary Figure 1. The PR proteins were selected by blasting with PR reported in the literature^111^. For the class of PR-1, results obtained here strongly suggest that some of them are pre-activated by *T. atroviride*-culture filtrate or *C. oleophila* cells pre-treatment (e. g. LOC111385823, LOC111385823, and LOC111384285); however, only the basic form of pathogenesis-related protein 1-like (LOC111384285) remained expressed in drupes only inoculated with *P. oleae* (treatment ID3) or when the pre-treatment with *C. oleophila* was also provided (treatment ID5) (Supplementary Figure 1). For β-1,3-glucanase (PR-2), the trend is more complex, being LOC111378392 the only one shared among the treatments and that could represent a threat to the target oomycete. The expression levels of the olive chitinases (PR-3, PR-4, PR-8, and PR-11) seem to have a better trend: the chitinase 10 (LOC111407547) and endochitinase EP3-like (LOC111393512) upregulation during the treatments with either *C. oleophila* or *T. atroviride*-culture filtrate seems to be a marker for the PR-3 weapons against the oomycete.

Among the PR-5 highly induced by *P. oleae* infection, the pathogenesis-related protein 5-like (LOC111397298) and the thaumatin-like protein (LOC111398526) were markedly differentially overexpressed (Supplementary Figure 1), but the pre-treatment with either *C. oleophila* or *T. atroviride*-culture filtrate slightly contributed to trigger the expression of the thaumatin-like protein (LOC111398526). The group of PR-6 includes important proteinase inhibitors, such as the cysteine protease inhibitors (relevant for blocking fungal cysteine proteases) and other enzyme inhibitors related to different classes of enzymes involved in the defense response^112,113^. Among these PR-6, as shown in Supplementary Figure 1, the polygalacturonase inhibitor LOC111382121 is the main PR-6 gene transcribed in response to fungal polygalacturonase. In this study, *C. oleophila* resulted to be an effective inducer of the expression of this gene, in contrast with *T. atroviride-*culture filtrate treatment. Neither *C. oleophila* or *T. atroviride*-culture filtrate could trigger the expression of the cysteine proteinase inhibitor B-like (LOC111390855), which was only induced by *P. oleae* infection itself. On the other hand, a set of inhibitors (proteinase inhibitor PSI-1.2-like LOC111390049 and LOC111373900, pectinesterase inhibitor 9-like LOC111377531, and kunitz trypsin inhibitor 2-like LOC111405411), including the cysteine proteinase inhibitor A-like (LOC111412304), were exclusively induced by *C. oleophila*.

Among the subtilisin-like protease (PR-7), serine protease belonging to the peptidases_S8 subfamily, which are already upregulated upon wounding alone, the highest upregulation was recorded for the gene encoding the subtilisin-like protease SBT1.9 (LOC111394370) in all the treatments. Previous studies demonstrated that the tomato P69 subtilase, which is a subtilisin, activates plant immunity since it can cut the pathogen-secreted apoplastic “small cysteine-rich secreted protein” PC2^114^. Downstream fragments are alerting the plant immunity by intracellular signaling. To evade the plant surveillance system, *Phytophthora* may have evolved protease inhibitors, such as the Kazal-like, protease inhibitors (EPIs) to block PC2 extracellular processing. Furthermore, *Phytophthora* may secrete cytoplasmic effectors to block the secretion of proteases (i.e., subtilisins and cysteine proteases) into the apoplast and, indirectly, block PC2-triggered immunity^115,116^. For PR-8, the acidic endochitinase-like (LOC111391829) was the main upregulated PR protein in response to *P. oleae* or *C. oleiphila* alone as compared the wound control. However, when both *P. oleae* and *C. oleophila* were present then it was downregulated. Other PR-8 expressed were hevamine-A-like (LOC111402962 and LOC111402963). The upregulation of PR-8 alone is hence not impressive to claim an active role in contrasting the infection.

The PR-9 proteins are lignin-forming peroxidases that fortify the cell wall by increasing the crosslinking of lignin. Results obtained here evidence that *P. oleae* infection determined the up-regulation of many PR-9 peroxidases (Supplementary Figure 1), but the treatment with either *C. oleophila* or *T. atroviride*-culture filtrate also contributed to increase the PR-9 expression level, especially for the lignin-forming anionic peroxidase-like (LOC111385717).

The PR-10 class is quite versatile in their biological action due to a loop that has small-chemical binding capabilities and diverse role in stress signaling^117^. In this study, the upregulation of some of them (e.g. pathogenesis-related protein STH-2-like -LOC111388404, LOC111388408, LOC111388411, and LOC111388413) is significative in olive drupes pre-treated with either *C. oleophila* or *T. atroviride*-culture filtrate as compared to olive drupes inoculated with *P. oleae* (Supplementary Figure 1). The highly expressed PR-11 are the acidic mammalian chitinases-like (LOC111382479, LOC111382480, and LOC111382484), but their expression is controversial, since wounding itself can trigger their higher expression and*T. atroviride* culture filtrate downregulated their expression.

Among the PR-12 class, the defensins-like protein 1 were not expressed. Some genes encoding PR-13 and PR-14 showed a slight upregulation in all the treatments.

The germin-like oxalate oxidases (PR-15) were also downregulated by *P. oleae* infection, even though treatment with either *C. oleophila* or *T. atroviride*-culture filtrate had the effect of upregulating some of them (Supplementary Figure 1).

Finally, the expression of PR-17 in olive drupes, which include important zinc apoplastic metalloproteinases, was relevant for the LOC111379582 and LOC111391744 in case of olive drupes only inoculated with *P. oleae* (treatment ID3) or drupes only pre-treated with *C. oleophila* (treatment ID7). Unfortunately, the combination of the pretreatment with either *C. oleophila* or *T. atroviride*-culture filtrate and *P. oleae* infection had the effect of reducing their expression (Supplementary Figure 1).

#### 2.2.6. Comparison 6: wounded olive drupes pre-treated with *C. oleophila* cell suspension (treatment ID7) versus wounded drupes pre-treated with sterile distilled water (treatment ID2)

In olive drupes, the treatment with *C. oleophila* (treatment ID7) triggered the differential upregulation of the lignin/lignan biosynthetic pathway, possibly to fortify the cell wall and produce anti-fungal compounds. Indeed, a lot of secoisolariciresinol dehydrogenase genes (LOC111412681, LOC111412273, LOC111412272, and LOC111412275) were overexpressed (**Supplementary Table 7)**. Two kunitz trypsin inhibitors (LOC111405411 and LOC111373900), two basic endochitinases (LOC111403475 and LOC111379188), and two pathogenesis-related proteins STH-like (LOC111388400 and LOC111388410) were highly upregulated as compared to the wound condition (treatment ID2). Meanwhile, the upregulation of metallothioneins (LOC111369652, LOC111375946, LOC111385831, and LOC111385129), thioredoxins (LOC111370790 and LOC111370789) and aquaporins (LOC111368643, LOC111408442, LOC111368644, and LOC111368645) might help the olive drupe to sustain a high level of ROS scavenging intracellularly and ROS export to the extracellular environment, such as superoxide (O_2_ ^−^), hydrogen peroxide (H O ) and the hydroxyl radical (•OH) ^118,119^.

#### 2.2.7. Comparison 7: wounded olive drupes pre-treated with *T. atroviride*-culture filtrate (treatment ID6) versus wounded drupes pre-treated with sterile distilled water (treatment ID2)

It is known that *Trichoderma* spp. produce metabolites able to trigger plant defense response against fungal pathogens^35,120,121^. In olive drupes pre-treated with *T. atroviride*-culture filtrate only 60 genes were upregulated and 176 downregulated (Supplementary Table 8). As for the treatment ID3 (*P. oleae* infection alone) also here the biosynthetic enzymes of the lignan biosynthesis was highly upregulated in particular the secoisolariciresinol dehydrogenase-like genes with L2FC≥3 (LOC111412681, LOC111412273, LOC111412272).In addition, polygalacturonase inhibitor (PR-6, LOC111385823), proteinase inhibitor PSI-1.2-like (LOC111390049), acidic glucan endo-1,3-beta-glucosidase (LOC111378392), kirola-like (PR-10, LOC111392831) and a basic endochitinase-like (LOC111379188) were also induced (Supplementary Table 8). In this study, it has been also found the induction of ethylene biosynthesis (1-aminocyclopropane-1-carboxylate oxidase-like LOC111370659). The activation of numerous disease-resistance-(Ethylene Responsive Factor, ERF, NAC, bHLH, and STK) and defense-response genes (DRP, ABC, and HSP) following treatment with *Trichoderma* spp. has been reported previously^122^. The NAC transcription factor 56-like (LOC111393458) and the protein DOWNY MILDEW RESISTANCE 6-like (LOC111393985) were also upregulated.

#### 2.2.8. Comparison 8: *P. oleae* differentially expressed genes during olive fruit infection (treatment ID3) as compared to its *in vitro* growth in several culture media

About 4513 genes were differentially expressed (2294 upregulated and 2219 downregulated) in *P. oleae* grown during the infection on olive fruits as compared to *P. oleae* grown in PDA, MEA, or Czapek media (Supplementary Table 9). A list of differentially expressed genes, identified as pathogen effectors, is given in Table 2. Most of the recorded effectors, related to the RxLR–dEER domain type, typical oomycete virulence proteins that should be translocated to the cytoplasm of the plant cell^123^. When secreted, RxLR proteins could be internalized by the binding with plant plasma membrane receptors^124^.Once internalized some RxLR proteins might interacts with resistance proteins (R proteins), and as a result this binding might trigger an HR and efficient callose deposition [i]. Crinkler (CRN) effectors play a significant role in the pathogenesis by manipulating host cell functions (*i.e.* cell death) to facilitate infection and suppress host immune responses^125^. Genes encoding CRN effectors were also abundant and highly upregulated (Table 2). Some of these effectors have been chosen for validation with RT-qPCR over the different time points (see Supplementary Table 15). Elicitins are the main pathogenic effectors in *Phytophthora* species^126^. Elicitin INF-1 represents the main important elicitin because it is known that induces a hypersensitive response (HR) and systemic acquired resistance (SAR)^127^. *P. oleae* possess two genes, g8411.t1 and g8407.t1, both predicted to encode the same proteins but with different length of leader peptide. A database of *P. oleae* proteins was made available as well as its first genome draft and the relative nucleotide and protein sequences at Zenodo (https://doi.org/10.5281/zenodo.11092245). *P. oleae* INF-1 were highly expressed in culture media but downregulated during olive fruit infection maybe to reduce HR and SAR and favor infection (Table 2). Most of the elicitins were downregulated except for a few that were upregulated (g1938.t1, g17483.t1, g18550.t1, g13418.t1, and g20317.t1) (Table 2).

**Table 2.**
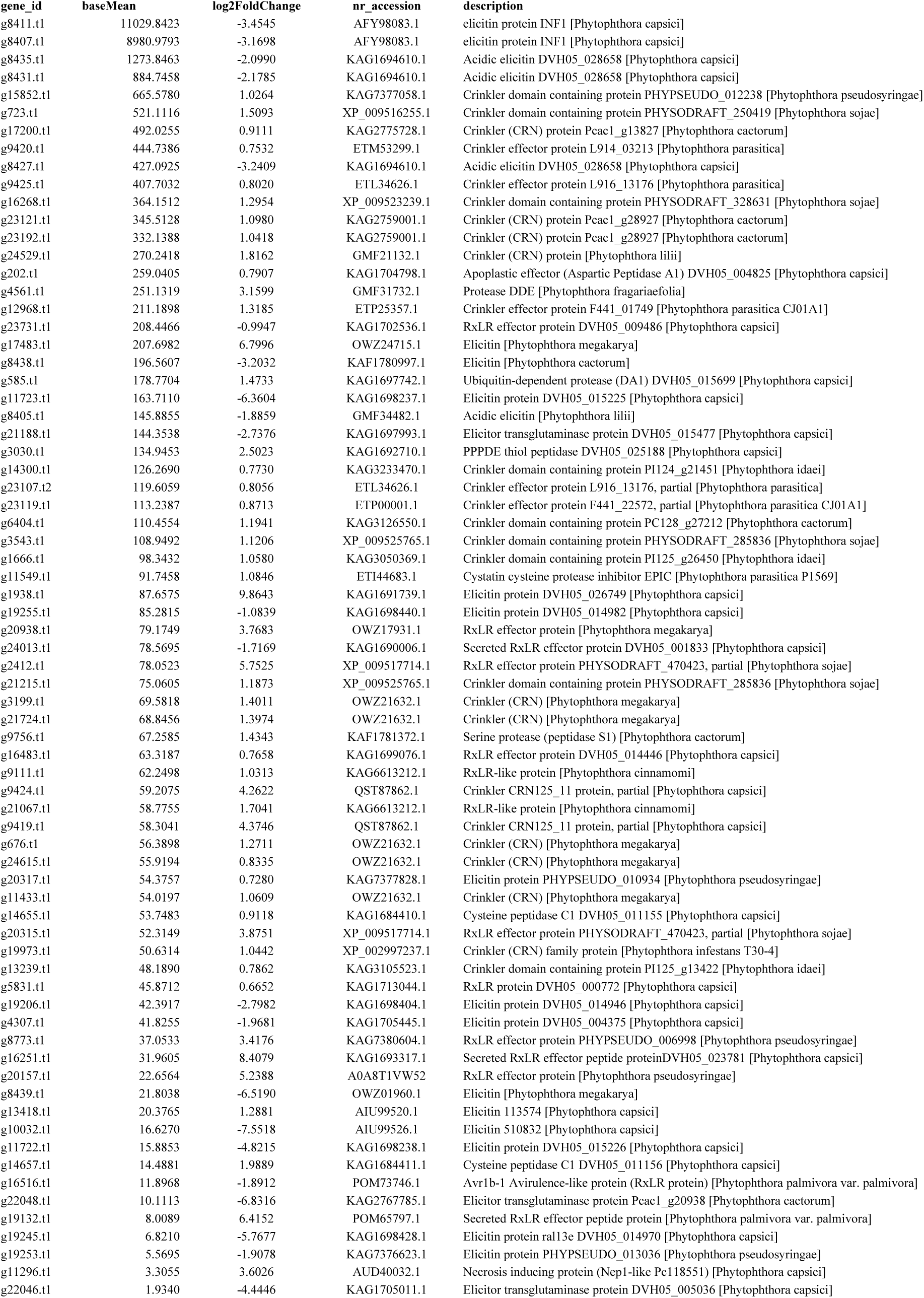
Selected upregulated mRNA related to P. oleae effectors differentially expressed into olive drupes during 72h of infection as compared to the mRNA expression during the growth in PDA (FDR =0.001). The gene expression has been filtered by the highest baseMean FPKM counts from Supplementary Table 11.

As for *P. oleae* proteases and protease inhibitors differentially expressed genes (statistically filtered; FDR=0.001), including cysteine proteases and serine protease types (Peptidase S10, serine carboxypeptidase, and chymotrypsin type), were downregulated. The cystatin cysteine protease inhibitor EPIC and the Kazal-like serine protease inhibitor were also downregulated (Supplementary Table 9). Necrosis inducing protein NPP1 or Nep-1 (g11296.t1) was upregulated while all the remaining homologues were downregulated (Supplementary Table 9).

As already mentioned, *Phytophthora* species. also secrete elicitins, whose function is to facilitate the onset of infection and its progress, because they are toxic proteins and most of the time induce necrotic and systemic hypersensitive response (HR) in the infected tissues. The interpretation of the less expressed elicitin protein INF1, known for its high toxicity, is that in certain cases INF-1 is responsible for the high virulence but it is also known to be highly expressed in artificial media as compared to biotroph growth in infected tissues^128^. Virulent races of fungal and bacterial pathogens are generally defined by their failure to express particular avirulence genes^129^. However, the loss or lack of elicitin production in *P. parasitica* was shown to increase its virulence on sensitive plants^130^. Here, the downregulation of *P. oleae* INF-1 (g8411.t1 and g8407.t1) and others (g8435.t1, g8431.t1, g8427.t1, g11723.t1, and g8405.t1) might instead have a strategic role to favor its virulence, since, for example, the *in vitro* elicitin production by *Phytophthora* spp. is strongly negatively correlated with their pathogenicity on tobacco^131^. The most upregulated elicitins, instead, were g8411.t1, g8407.t1, g1938.t1, g17483.t1, and g18550.t1 (Supplementary Table 9). Other avirulent effectors were also highly upregulated, like cysteine peptidase C1 domain containing protein (g14657.t1) and cystatin cysteine protease inhibitor EPIC-like (g11549.t1) (Table 2).

#### 2.2.9. Comparison 9: differentially expressed genes of *P. oleae* during olive drupe infection (treatment ID3) as compared to olive drupe inoculated with *P. oleae* and pre-treated with either *C. oleophila* (treatment ID5) or *T. atroviride*-culture filtrate (treatment ID4)

Supplementary Table 10 shows 3716 (2116 upregulated and 1600 downregulated) and differentially expressed mRNAs related to *P. oleae* during pre-treatment of olive drupes with either *C. oleophila* (treatment ID5) or *T. atroviride*-culture filtrate (treatment ID4) versus olive drupes infected with *P. oleae* (treatment ID3), 72 hpi (FDR =0.001). Supplementary Tables 11 and 12 show the *P. oleae* differential mRNA expression in the comparisons ‘olive drupes pre-treated with *C. oleophila* and inoculated with *P. oleae* (treatment ID5)’ versus ‘olive drupes only inoculated with *P. oleae* (treatment ID3)’, and ‘olive drupes pre-treated with *T. atroviride*-culture filtrate and inoculated with *P. oleae* (treatment ID4)’ versus ‘olive drupes only inoculated with *P. oleae* (treatment ID3)’, respectively.

In the pre-treatment with *T. atroviride*-culture filtrate, the gene-encoding for the elicitin proteins INF-1 (g8411.t1 and g8407.t1) have been slightly upregulated as compared to drupes only inoculated with *P. oleae*; however, *P. oleae*-inoculated drupes pre-treated with *C. oleophila* showed a slight downregulation (L2FC= -0.8) (Supplementary Table 11). The differentially expressed elicitin genes that were not down-regulated in *P. oleae*-inoculated olive drupes pre-treated with either *C. oleophila* or *T. atroviride*-culture filtrate were g21198.t1, g10684.t1, g17483.t1, g24094.t1, and g4307.t1 (Supplementary Tables 10, 11); other genes-encoding for elicitins were downregulated by pre-treatment with either *C. oleophila* or *T. atroviride*-culture filtrate (e. g. g13418.t1, g1938.t1, g19255.t1).

To evaluate the efficacy of the antagonistic effect of the treatment with either *C. oleophila* or *T. atroviride*-culture filtrate versus the infection of *P. oleae* alone and, for comparison purposes, genes, vs. axenic cultures of *P. oleae* grown in PDA, MEA and CZ (CZAPEK) the DESeq2 results at 72 hpi were analyzed using the Principal Component Analysis (PCA)(Figure 7). The sum of all these components (PC) accounted for 76% of the total variance, while PC1 and PC2 represented 53% and 23%, respectively. Although PCA analysis is usually significant when the percentage of variance in the first three components is at least 80%, it may be used to find correlations based on the co-variance matrix. Figure 7 shows that four distinct groups were grouped apart. Specifically, olive drupes inoculated with *P. oleae* were clearly separated from *P. oleae-* inoculated drupes pre-treated with either *C. oleophila* or *T. atroviride*-culture filtrate, as well as from the axenic cultures of *P. oleae* on CZ and MEA media. Only the culture of *Phytophthora oleae* on PDA medium is singularly grouped in the quadrant with negative values on the abscissa and ordinate.

**Figure 7.**
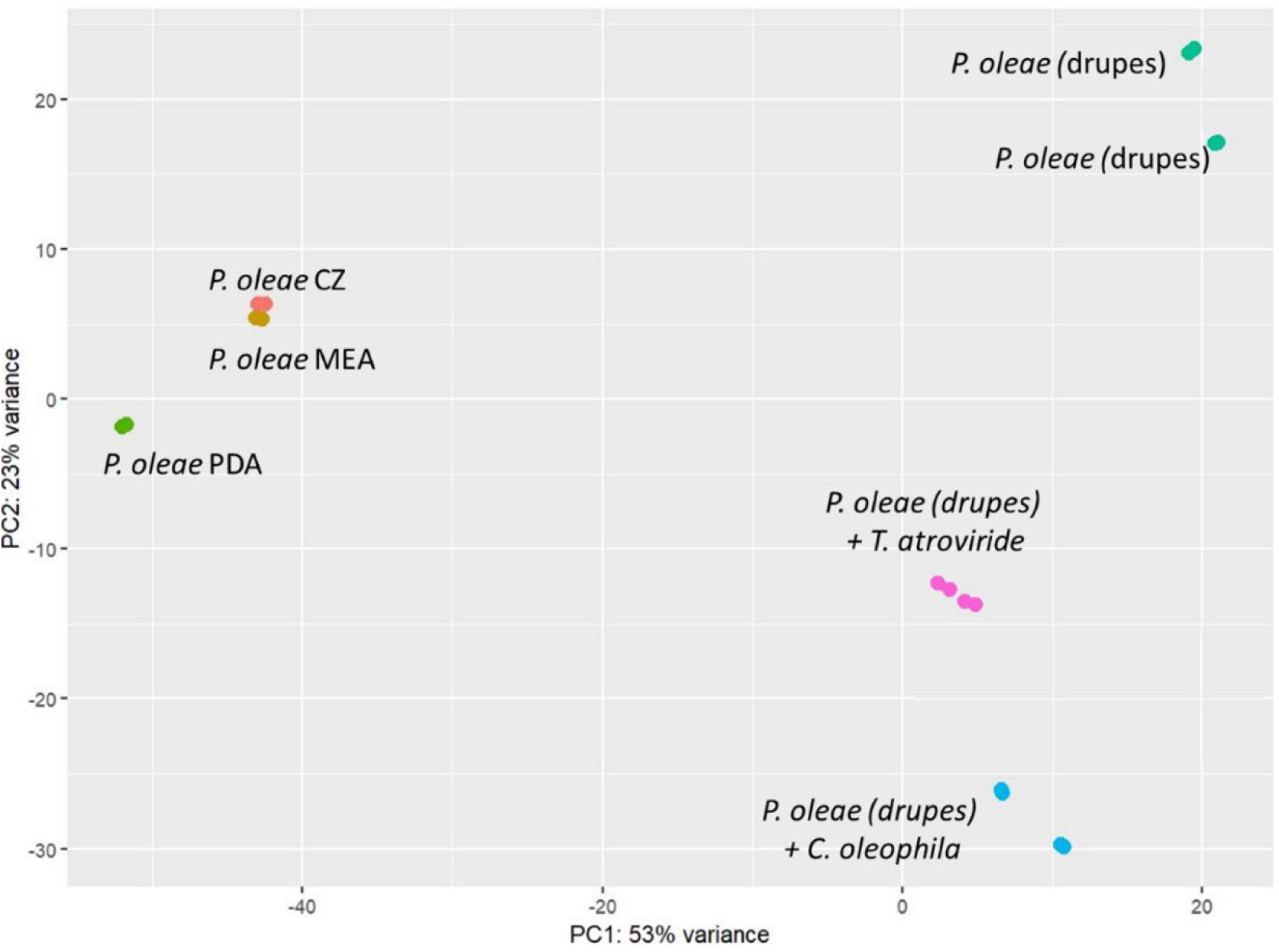
PCA plot including all the *P. oleae* samples alone or with either *C. oleophila* or *T. atroviride*-culture filtrate from 72hpi DESEq2 data. The horizontal axis (PC1) represents the distance from *P. oleae* grown in artificial media (CZ, MEA, or PDA) proportional to the variance of pathogenicity (rAUDPC) starting from the coordinates 0, 0. Negative PC1 variance means no infection. The closer the horizontal distance from 0, 0, the less is the degree of infectivity for the cases where infection is ongoing. The vertical variance of this PCA describes the variation of genes involved in severity of the disease as compared to original characteristics (no infection) of *P. oleae* grown in artificial media (CZ, MEA, or PDA). The distance/variance in PC2 between *P. oleae* infecting the drupe alone has a positive value while the distance/variance in PC2 of *P. oleae* infecting the drupe plus either *C. oleophila* or *T. atroviride*-culture filtrate is proportional to the reduction of infectivity detected in each mixed condition (*P. oleae* + either *C. oleophila* or *T. atroviride*-culture filtrate variance) where was observed a negative variance in PC2.

Furthermore, several cluster analyses have been performed to evaluate the changes in genes-encoding for a number of pathogenic enzymes and effectors. To this aim, elicitins, crinkler, peptidases, and RxLR effectors LF2C from DESeq2 data expression were chosen (Figure 8). Some examples include peptidases as the cysteine peptidase C1 (g9173.t1), which are downregulated when *P. oleae* is inoculated in olive drupes, but upregulated when either *C. oleophila* or *T. atroviride*-culture filtrate are included as pre-treatment. This could mean that perhaps the cysteine peptidase C1 (g9173.t1) could serve as weapon against other surrounding fungi. An opposite trend was observed for the Peptidase s59 (g4376.t1) (Figure 8A) and for the RxLR effectors (Figure 8C). The genes-encoding for the RxLR effector proteins g6343.t1, g19400.t1, g16251.t1, g19132.t1, g24597.t1, g7454.t1, and g6774.t1, for example, were clearly upregulated in *P. oleae*-inoculated and non-treated olive drupes (treatment ID3) but downregulated when either *C. oleophila* (ID5) or *T. atroviride*-culture filtrate (ID4) where provided as pre-treatment (Figure 8C). Most of those highly expressed *P. oleae* RxLR effectors might attenuates the responses to SA like including of PR1 as in the case of *Arabidopsis* infected with *Hyaloperonospora arabidopsidis* (Hpa Emoy2)^132^. In facts, g6343.t1, and g7454.t1 are homologous respectively to HaRxLL465b (BAP69127.1) and HaRxL24b (CCC55787.1) highly expressed in Hpa Emoy2 [p]. Besides, *P. oleae* RxLR effector g19132.t1 possesses also one WY motifs, known for being necessary for the infection and RNA silencing suppression activity^133^.

**Figure 8.**
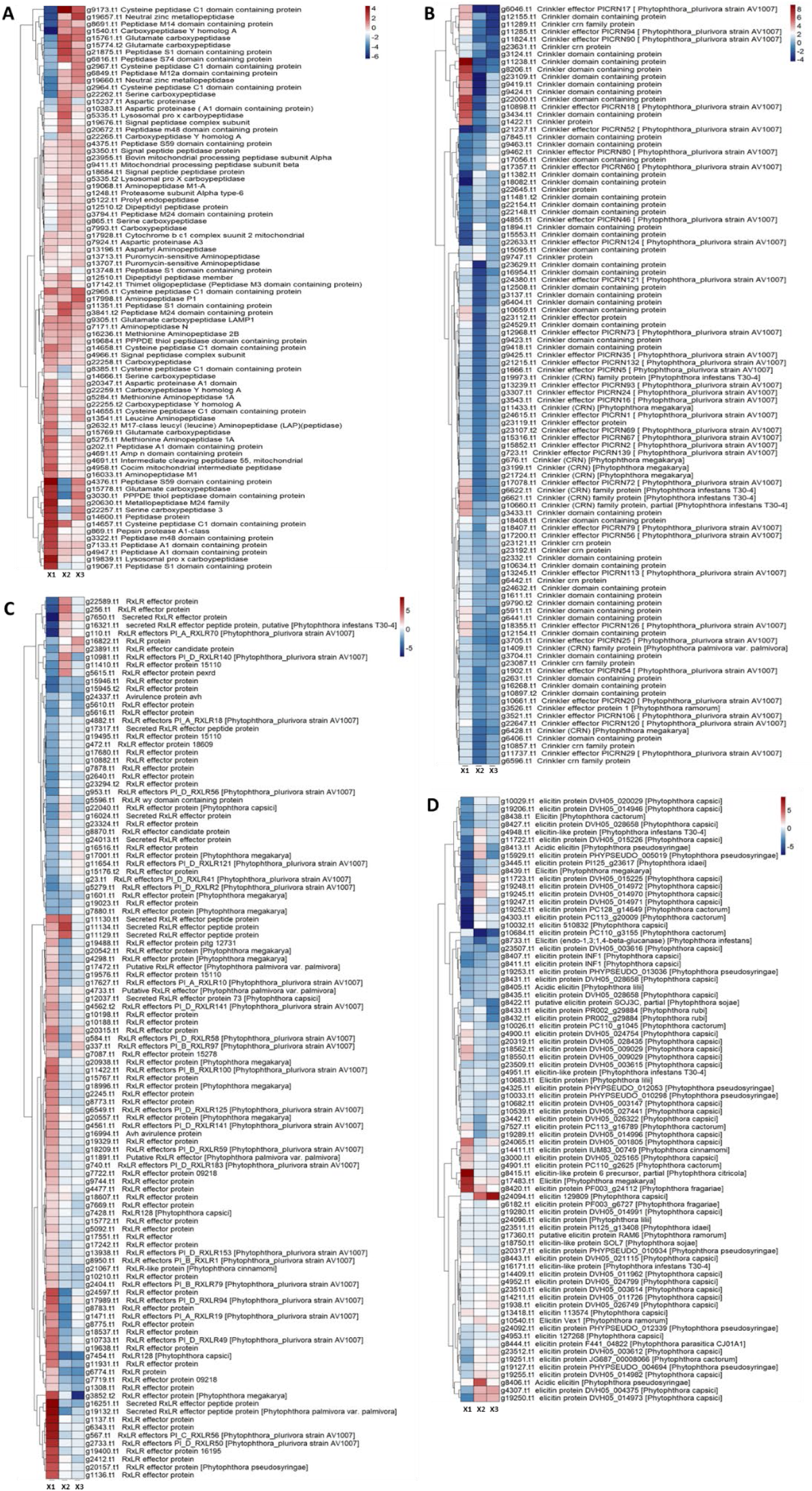
Cluster heatmap analyses of *P. oleae* peptidases, Crinkler, RxLR effectors and elicitins differentially expressed in three conditions tested. X1= L2FC of *P. oleae* grown on olive drupes vs. *P. oleae* grown on artificial media; X2= L2FC of *P. oleae* grown on drupes plus *C. oleophila* vs. *P. oleae* grown on drupes; X3= L2FC of *P. oleae* grown on drupes plus *T. atroviride* filtrate vs. *P. oleae* grown on drupes.

The trend of the *P. oleae* CRN78 homologous isogenes (g11433.t1, g676.t1, g3199.t1, g21724.t1, g24615.t1) was characterized by an up-regulation when *P. oleae* was inoculated in non-pre-treated olive drupes (L2FC > 1.2) (ID3) and by a downregulation in drupes pre-treated with either *C. oleophila* (L2FC<-1.2) (ID5) or *T. atroviride* (L2FC<0.4) (ID4) (Figure 6B). The Crinkler effector CRN78 of *P. soja* (XP_009521873.1) is known to enhance oomycete pathogen infection by inhibiting plant immunity through a unique mechanism^134^.

*P. sojae* effector CRN78 contains a secretion signal peptide (SP), a typical LXLFLAK and HVLVVVP motifs at its N terminal; this latter is predicted to be a serine/threonine kinase domain. It has been demonstrated that CRN78 mediates phosphorylation of NbPIP2;2 at Ser279 and induces its degradation via a 26S-dependent pathway^134^. Since the aquaporin PIP2;2 acts as a H2O2 transporter into the apoplast, its degradation substantially decreases extracellular ROS.

#### 2.2.10. RT-qPCR validation of selected genes of *Olea europaea* and *Phytophthora oleae*.

A selection of differentially expressed genes of olive drupes (Supplementary Table 15), as analyzed by RNAseq at 72 hpi, have been validated during the entire time course considered (24, 72, and 168 hpi). Figure 9 shows the trend of expression of those genes. At 24 hpi, the gene-encoding for the pathogenesis-related protein STH-2-like (XM_023013103.1, LOC111388413) (Figure 9A) was significantly up-regulated in all the treatments, but in wounded drupes; thus, indicating that this gene is positively transcribed as a consequence of biotic stresses. Furthermore, *P. oleae* inoculated drupes pre-treated with either *T. atroviride*-culture filtrate or *C. oleophila* showed the highest upregulation values. A similar trend was also observed at 72 hpi, although less marked in olive drupes only inoculated with *P. oleae* (ID3) and those also pre-treated with *T. atroviride*-culture filtrate (ID4). Finally, at 168 hpi, the most significant up-regulation of the gene-encoding for the pathogenesis-related protein STH-2-like was recorded in *P. oleae*-inoculated olive drupes pre-treated with *T. atroviride*-culture filtrate. Overall,, throughout all the infection period considered, it is constant the higher upregulation of this PR protein in the presence of both *C. oleophila* and *T. atroviride* culture filtrate, phenomenon already known for *T. harzianum* in case of sunflower challenged by *Rhizoctonia solani*^135^. The gene-encoding for the BON1-associated protein 2-like (LOC111391572, XM_023016810.1) (Figure 9B) showed a marked upregulation in drupes only inoculated with *P. oleae* (ID03) at the three time points of measurement. BON1 is a calcium-dependent phospholipid-binding protein interacting with the leucine-rich-repeat receptor-like kinases BIR1 (BAK1-interacting receptor-like kinase 1) and pathogen-associated molecular pattern (PAMP) receptor regulator BAK1^136^. BON1-associated protein participates to the protein-protein regulation of the PAMP signaling of which BAK1 represent a hub of signals functioning as negative regulator role related to cell death and defense responses^137^. Results obtained in this study suggest that the infection by *P. oleae* caused the upregulation of BON1-associated protein, probably suppressing both cell death and reducing the PAMP response in activating the defense response. Notably, all the treatments that included either *C. oleophila* or *T. atroviride*-culture filtrate showed a generalized low up-regulation or a down-regulation. This study also analyzed the trend of the gene-encoding for FERONIA (Figure 9C), an LRK-receptor-like protein kinase (LOC111374777, XM_022997515.1), involved in the plant immunity by inhibiting JA signaling^89^. Hence, the transcription of this gene is supposed to exert a negative function toward *P. oleae* pathogenicity. Results obtained here show that at 24 hpi the FERONIA encoding-gene was significantly up-regulated in *P. oleae*-inoculated olive drupes pre-treated with *T. atroviride*-culture filtrate (ID4) and in olive drupes only pre-treated with *C. oleophila* (ID5); at 72 hpi the modulation in the transcription of the gene was significant just in *P. oleae*-inoculated olive drupes pre-treated with *T. atroviride*-culture filtrate (ID4); at 168 hpi, no-significant differential transcriptions were recorded in drupes from any treatment.

**Figure 9.**
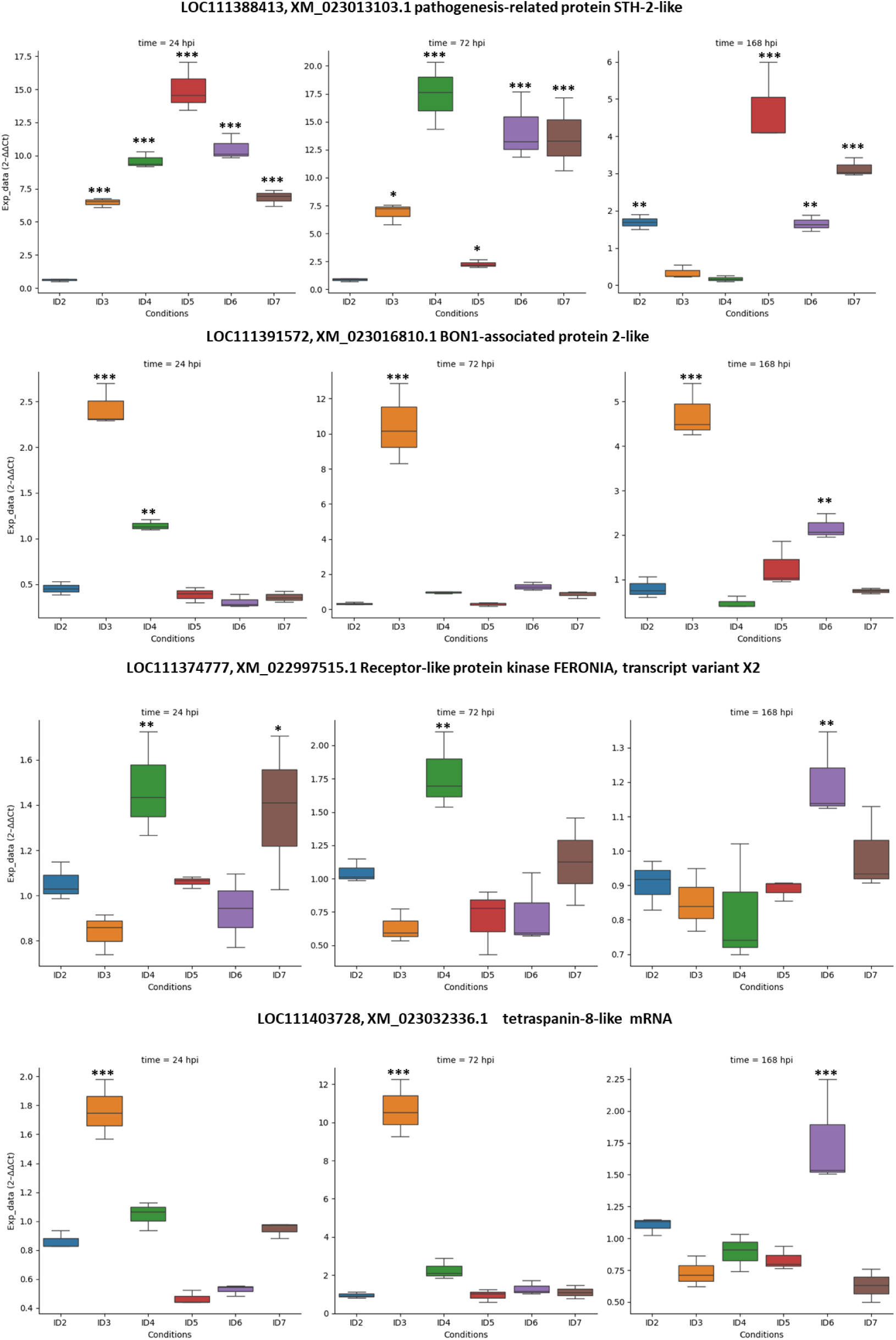
RT-qPCR validation of chosen DEGs for *Olea europaea* at three time points and under different conditions. Asterisks represents statistically different samples according to Dunnett’s test (∗ = p < 0.05, ∗∗ = p < 0.01, and ∗∗∗ = p < 0.001), as compared to unwounded fruits. The normalization has been performed according to the tubulin alpha-3 chain (LOC111371391, XM_022993342.1). The RT-qPCR primers are shown in Supplementary Table 15. The bars represent the standard deviation (SD). Conditions legend: ID2, wound; ID3, wound + *P. oleae*; ID4, wound + *T. atroviride* + *P. oleae*; ID5, wound + *C. oleophila* + *P. oleae*; ID6, wound + *T. atroviride*; ID7, wound + *C. oleophila*.

As mentioned, the plant small extracellular vesicles (EVs) or exosomes (EXOs) that carry plant defense enzymes, PRs and the ROS generating machine, were supposed accumulating toward the pathogen entrance hot spot. Plant exosomes contain DADPH oxidases, PM-ATPases, and tetraspanins that mediate PM-fusion (unloading of content). Many tetraspanins are involved in EXOs trafficking and here, the gene-encoding for the tetraspanin-8-like (LOC111403728, XM_023032336.1) was chosen as a top differentially expressed gene for RT-qPCR-analysis (Figure 9D). At the time points 24- and 72-hpi, the significant upregulation of the tetraspanin-8-like in drupes only inoculated with *P. oleae* (ID3), was interpreted as the attempt of plant cells to counteract the oomycete pathogen by ROS and PRs. Conversely, the downregulation in *P. oleae* inoculated drupes pre-treated with either *C. oleophila* (ID5) or *T. atroviride*-culture filtrate (ID4), although non-significant, suggests that the presence of either *C. oleophila* or *T. atroviride*-culture filtrate could have downregulated the ROS and PRs mediated mechanism by their own extracellular enzymes/effectors to help the plant immunity by by-passing the exosomes mechanism of defense. This interpretation of the results is supported by the fact that at 168-hpi, drupes only inoculated with *P. oleae* (ID3) exhibited a downregulation of tetraspanin-8-like gene, indicating that the advanced stage of infection could have completely compromised any plant defense response. At this time point, the only significant up-regulation was recorded in olive drupes treated with *T. atroviride*-culture filtrate.

In this study, the RT-qPCR validation of some selected *P. oleae* differentially expressed encoding effectors (Supplementary Table 15) at all time points considered was also carried out. As comparison, the mean value of expression level of the *P. oleae* genes grown in artificial media (PDA, MEA, or Czapek) was used.

The Crinkler (CRN) gene g723.t1 (TRINITY_DN5906_c2_g5_i1) (Figure 10A) is homologous of the Crinkler effector of *P. soja* (XP_009516255.1) and the Crinkler effector protein 5 of *P. infestans* (Q2M408.1) The CRN proteins are supposed to be secreted by the oomycete into the host apoplast and, by an unknown mechanism, they can penetrate and exert their toxic necrotic function against the host. Results of this study strongly suggest that the presence of either *C. oleophila* (ID5) or *T. atroviride*-culture filtrate (ID4) smoothed the upregulation of the Crinkler (CRN) gene g723.t1, as it is markedly upregulated in the treatment where *P. oleae* is alone (ID3) (Figure 10A). A similar trend was recorded for the Elicitin g17483.t1 (Figure 10B). The expression trend of the INF-1 elicitin encoding-genes (g8411.t1 and g8407.t1) (Figure 10C) aligns with the RNAseq analysis findings at 72 hpi. In the treatment where *P. oleae* is alone (ID3), the INF-1 elicitin is down-regulated at 24 hpi. Additionally, at 72 and 168 hpi, its upregulation was significantly lower than in drupes that also included pre-treatment with either *C. oleophila* (ID5) or *T. atroviride*-culture filtrate (ID4). This INF-1 elicitin gene trend in treatment ID3, which overall exhibited consistently lower positive regulation compared to the other two treatments, might be attributed to the potential establishment of an ’infection barrier’ by the presence of either *C. oleophila* or *T. atroviride*-culture filtrate. Hence, to overcome this barrier, *P. oleae* might have increased the synthesis of INF-1 elicitin.

**Figure 10.**
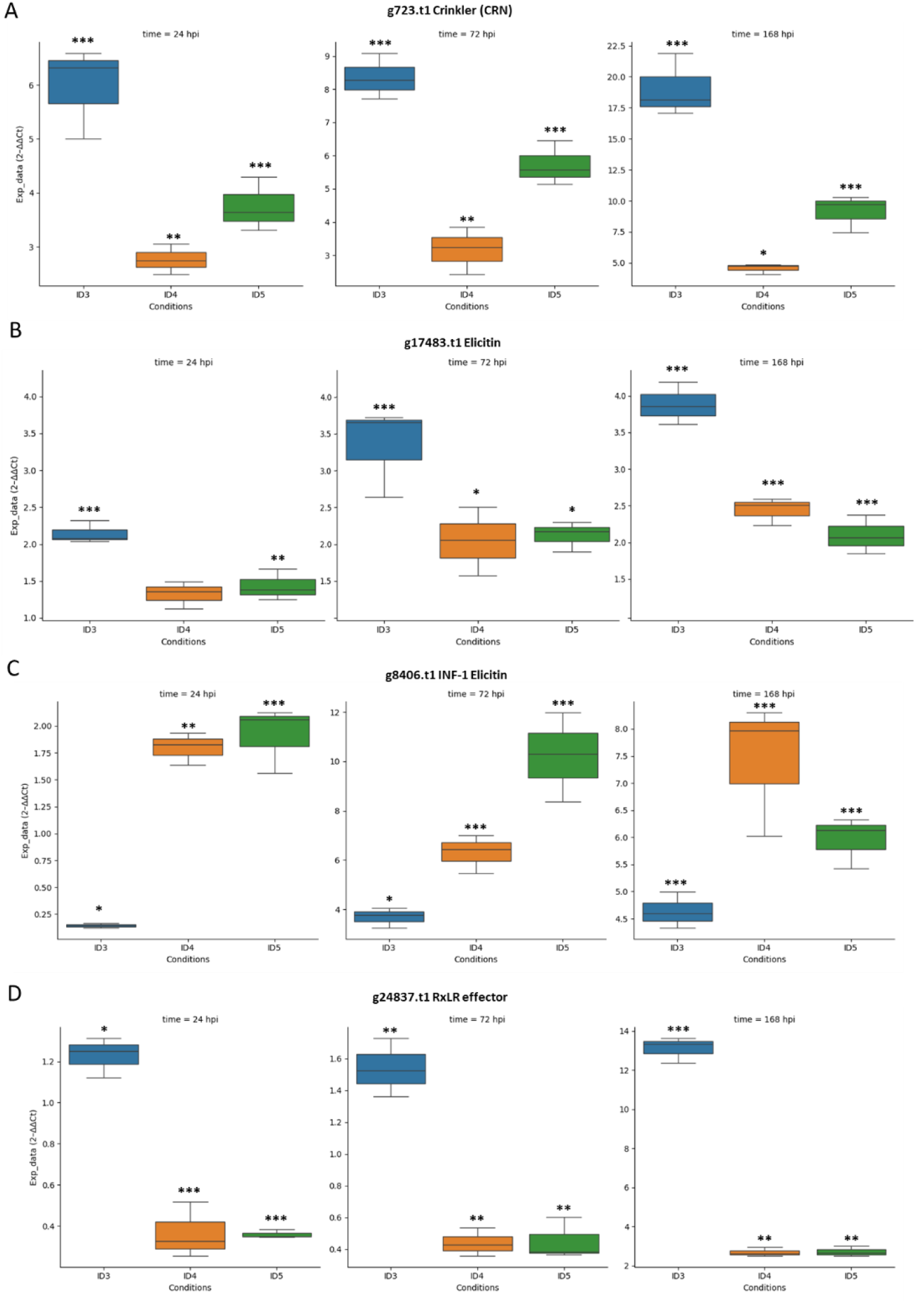
RT-qPCR validation of chosen DEGs for *P. oleae* during the three time points and different conditions. Asterisks represent statistically different responses as compared to the expression in artificial media, according to Dunnett’s test (∗ = p < 0.05, ∗∗ = p < 0.01, and ∗∗∗ = p < 0.001). The normalization has been performed according to the tubulin beta chain g8215.t2 (TRINITY_DN6095_c0_g1_i1). The RT-qPCR primers are shown in Supplementary Table 15. The bars represent the standard deviation (SD).

Another host-translocated effector belonging to the RxLR family, the *P. oleae* g24837.t1 (TRINITY_DN8100_c0_g1_i1) (Figure 10D) homologous to HaRxLL438 (BAP69100.1) of *H. arabidopsidis* Emoy2, upregulated upon infection, has also been detected as highly expressed in the case of infection with *P. oleae* alone, but downregulated over time following treatments with either *C. oleophila* or *T. atroviride*-culture filtrate.

## 3. Discussion

This study demonstrated the yeast *C. oleophila* and the culture filtrate of the filamentous fungus *T. atroviride* were effective in reducing the severity of symptoms of soft rot in olive drupes infected by *P. oleae*. To the best of our knowledge, this is the first report of *in planta* efficacy of these treatments, a BCA and a culture filtrate of an antagonistic fungus. Results are promising, also in the light of the transcriptome analysis, indicating the activation of plant immune system by the yeast and culture filtrate tested in this study. This would suggest the possibility that both *C. oleophila* and *T. atroviride* culture filtrate can be effective against a broad spectrum of fruit diseases caused by necrotrophic pathogens. Indeed, *C. oleophila* strain O is already available as a commercial bioproduct registered for treatments of post-harvest fungal rots of several fruit crops, while the practical application of *Trichoderma atroviride* metabolites as natural substances for the biocontrol of plant diseases is almost unexplored. Moreover, the pathosystem *P. oleae*/olive drupe treated with either *Trichoderma* culture filtrate or *C. oleophila* cell suspension proved to be an interesting model to analyze the gene expression in a multiple-actor (pathogen/host plant/treatment) interaction. The transcriptome analysis of this interaction provided a better understanding of the defense responses of olive to stressors, including the abiotic stress provoked by wound inoculation, and the mechanisms underlying the biocidal activity of two very diverse means. The components of the model were chosen and assembled in various combinations based on a rationale. *Phytophthora oleae* is an emergent olive pathogen and the genus *Phytophthora* includes numerous species that are responsible of serious diseases of crop, forest, ornamental and landscape plants. ‘Coratina’, originating from the Apulia region (southern Italy), is one of the most popular Italian olive varieties for oil production. It is an ancient variety thought to express many wild resistance genes and to be closer to *O. europaea* var. *sylvestris* (taxid:158386), the wild relative of *O. europaea* var. *sativa*, in genomic sequence. Hence, RNAseq analysis has been performed using the O_europaea_v1 genome assembly (GCF_002742605.1). As in many studies it was demonstrated the strong efficacy of different strains of *C. oleophila* against post-harvest fungal diseases of edible fruit crops such as apples, bananas, citrus and strawberries^93,138^, this BCA was selected for the treatment of olive drupes. The choice of evaluating *T. atroviride* culture filtrate as an eco-friendly treatment was suggested by the high efficient production of bioactive secondary metabolites in liquid culture with antifungal activity by other *Trichoderma* spp. ^22,24,139–1^. ^141^. The elicitation of the plant immune system by liquid culture filtrate of *T. atroviride*, in this study, has perhaps been triggered by the elicitin Epl1 (XP_013937770.1) Cerato-platanin, which was found the most highly expressed protein by proteomics analysis (Table 1). Ep1 potential in triggering plant defense responses is well known^107,108^.

The time course of olive rot severity for each treatment, as represented by rAUDPC, is in somewhat in agreement with the PCA plot (Figure 7), which shows the variance of the expression of genes involved in pathogenicity as compared to the *P. oleae* growth in artificial medium (baseline or zero variance) or during infection (positive variance) or infection and pre-treatment with either *C. oleophila* or *T. atroviride*-culture filtrate (negative variance). In both cases, pre-treatment with *T. atroviride*-culture filtrate had the effect of attenuating the virulence of *P. oleae*. *P. oleae* effectors had a marked downregulation as a consequence of pre- treatment with *T. atroviride*-culture filtrate. This down regulation was probably due to a combined effect of the plant tissue pre-elicited by *T. atroviride* elicitin (e.g. Epl1 Cerato-platanin) and other bioactive metabolites of the culture filtrate.

The RNAseq analysis of olive gene differentially expressed during the infection by *P. oleae* showed the most upregulated genes were EXORDIUM-like isogenes, known to inhibit cell proliferation^52^ and which in soybean are associated with resistance to *P. sojae*^54^. In inoculated olives the ethylene biosynthesis occurred very early during the infection process and was triggered by wounding itself, as revealed by the expression of 1-aminocyclopropane-1-carboxylate synthase-like genes (ACC synthases; LOC111390291, LOC111392037 and LOC111370655) (Figure 2; Supplementary Table 1). In olive drupes only inoculated with *P. oleae*, 1-aminocyclopropane-1-carboxylate oxidase 1-like (ACO, LOC111366119) was also highly upregulated (Figure 3; Supplementary Table 2) and represented the last step in ET biosynthesis. Infection by *P. oleae* upregulated significantly also Ca^2+^ signaling, more than wounding itself as shown by the upregulation of protein SRC2 homolog (LOC111386812; Figure 3), which is the target of the main *P. oleae* elicitin INF-1 (g8411.t1 and g8407.t1).

The most upregulated gene in olive fruit during *P. oleae* infection, the BON1-associated protein 2 (LOC111391572), function as a negative regulator of cell death and defense responses in association with BAP1 and negatively regulating the R-protein “suppressor of npr1-1, constitutive 1” (SNC1)^56,57^. Infection by *P. oleae* led to suppression of the natural immunity response by regulating BON1/SNC1 at the site of infection, but perhaps was not sufficient to inhibit the production of salicylic acid and pipecolic acid by maintaining the upregulated expression of SAR DEFICIENT 1-like (LOC111402349 and LOC111377821), initiating the systemic acquired resistance (SAR) in distal parts of the plant^60,62^. As already reminded, SA and Pip trigger the activation of immune-related genes, such as many pathogenesis-related protein genes (PRs), including pathogenesis-related proteins STH-21-like, kirola-like proteins, hevamine-A-like, thaumatin-like proteins, cysteine proteinase inhibitors A/B-like, kunitz trypsin inhibitor 2-like, chitinases, endoglucanases, and β-1,3-glucanases. These are upregulated during *P. oleae* infection (Figure 3; Supplementary Table 2) and significantly even more following treatment with *C. oleophila* (Table 1; Supplementary Table 3) and *T. atroviride*-culture filtrate (Supplementary Table 4).

The receptor kinase FERONIA is downregulated by *P. oleae* infection and upregulated by the presence of *C. oleophila* and *T. atroviride* extracts as also validated over time by qPCR(Figure 9). During cell elongation, RALF23 plant peptides, when cleaved by the site-1 protease (S1P, a serine protease of the family of proprotein convertase subtilisin/kexins), can create a complex with FER and the binding activate it kinase activity resulting in phosphorylation of plasma membrane H^+^-ATPase 2 at serine-899, causing inhibition of proton transport and in general an arrest of root growth^142^. The FER activation suppresses also flg22-induced reactive oxygen species (ROS) burst and immune responses. This suppression is achieved by destabilizing the complex formation of EFR and FLS2 with their co-receptor BAK1, illustrating how RALF-FER signaling can modulate immune signaling pathways^143^. Pathogens, such as fungi, have evolved strategies to manipulate plant signaling pathways to facilitate infection. The discovery of potential RALF peptides in fungal and bacterial genomes suggests that pathogens might exploit RALF-FER signaling to manipulate plant immune responses and facilitate infection. Some fungal pathogens (e. g. *Fusarium oxysporum*) produce peptides that mimic plant signaling molecules, known as RALFs-like peptides, which can interact with FER by activating it and hence promote infection. These pathogen-derived RALF mimics can hijack the FER signaling pathway, leading to increased virulence and successful colonization of the plant host. The interaction between RALF peptides and FER triggers a signaling cascade that can inhibit cell growth, as seen in the inhibition of primary root growth in *Arabidopsis* upon binding of RALF1 to FER^144^. For instance, *F. oxysporum f. sp. lycopersici* RALF-like peptides inhibited the growth of tomato seedlings and elicited responses in tomato and *Nicotiana benthamiana* typical of endogenous plant RALF peptides (reactive oxygen species burst, induced alkalinization and mitogen-activated protein kinase activation)^145^. To make the hypothesis that FER is upregulated due to its inhibition by *P. oleae* effectors, we searched into a group of small proteins, top differentially expressed, in case of *P. oleae* infection (Supplementary Table 9) and aligned their protein sequence to known RALF peptides from *O. europaea* and/or fungal RALF-like peptide homologues (Supplementary Figure 2). The alignment revealed a conservation of critical cysteine residues and FER binding motifs even if those were more similar to the *Colletotrichum higginsianum* RALF-like protein (CCF44719) rather than *F. graminearum* (FG05_30327). *P. oleae* putative RALF-like homologues, differentially expressed, were: g3545.t1, g8557.t1 and g5663.t1. The hypothesis here is that *P. oleae* putative RALF-like proteins/peptides might activate FERONIA, and hence *P. oleae* could block the JA, COR and SA signaling, facilitating its pathogenesis. We need, of course, to confirm by other experiments whether the putative bioinformatics identified *P. oleae* RALF-like proteins here (Supplementary Figure 2) really interact with FERONIA.

Many R-genes were upregulated during *P. oleae* infection (Figure 3; Supplementary Table 2), probably acting in conjunction with WRKY TFs and F-box/LRR-repeat proteins to express such PRs. During 72hpi of *T. atroviride*-culture filtrate pre-treatment followed by *P. oleae* infection, the R-genes were particularly upregulated: LRR receptor-like serine/threonine-protein kinase At1g06840 (LOC111371641), LEAF RUST 10 DISEASE-RESISTANCE LOCUS RECEPTOR-LIKE PROTEIN KINASE-like 1.1 (LOC111390435), MLO-like protein 2 (LOC111370597), rust resistance kinase Lr10-like (LOC111411655), pto-interacting protein 1-like (LOC111410523), WRKY transcription factor 20 (LOC111391535), and the LRR receptor-like serine/threonine-protein kinase At1g06840 (LOC111393227). Calcium (Ca^2+^) signaling plays a pivotal role in plant immunity, functioning as a key secondary messenger in response to various stimuli, including pathogen attack. In plants, Ca^2+^ signaling is central to both pattern-triggered immunity (PTI) and effector-triggered immunity (ETI), with the generation of characteristic cytoplasmic Ca^2+^ elevations being common to both^146^. The interaction between FERONIA and MLO-like proteins (e.g. NORTIA, NTA) is known in pollen tube reception and fungal invasion causing a release of intracellular Ca^2+^ and related signaling^91^. Here, FERONIA and MLO-like protein2 might have interacted and triggered the same Ca^2+^ release and signaling followed by upregulation of many Ca^2+^ binding proteins for further signaling and immunity activation. On another hand, *T. atroviride-*culture filtrate pre-treatment triggered the upregulated of other R-genes: the LRR receptor-like serine/threonine-protein kinase At1g06840 (LOC111371641), LRR receptor-like serine/threonine-protein kinase At1g06840 (LOC111393227), and the MLO-like protein 8 (LOC111401243). Besides, BON1-associated protein 2 (LOC111391572) was downregulated following pre-treatment with either *C. oleophila* or *T. atroviride*-culture filtrate (Supplementary Tables 3, and 4).

Necrotrophic pathogens and beneficial antagonist microorganisms induce or prime ET- and JA-dependent signaling pathways. For example, the necrotrophic pathogen *Botrytis cinerea* causes a rapid activation of ET biosynthesis in *Arabidopsis* by the action of Mitogen Activated Protein Kinase3 (MAPK3) and MAPK6 that phosphorylate the ET biosynthesis proteins 1-Aminocyclopropane-1-Carboxilic Acid (ACC) Synthase 2 (ACS2) and ACS6^147^. In the case of *P. oleae* infection alone, ethylene biosynthesis was highly upregulated since it was found that the final step involving biosynthetic enzyme 1-aminocyclopropane-1-carboxylate oxidase 1-like (ACO, LOC111366119), was very highly upregulated as compared to wounded fruit (L2FC≥ 7.6; Figure 3; Supplementary Table 2). Besides the boost to ET biosynthesis, it was observed a significant increase in JA biosynthesis and an upregulation of many ET responsive genes. The Apetala 2 (AP2)/Ethylene Response Factor (ERF) seems to be the driver of the ET/JA-mediated defense pathway. Its homologue in olive is the ethylene-responsive TF 1B-like (ERF-1B) of which many isogenes are present in the genome. Upon *P. oleae* infection, many ERF-1B like factors were upregulated (Supplementary Table 2). As described above, JA binds to the JA receptor Coronatine Insensitive 1 (COI1, LOC111390662 in *O. europeae*), which results in the degradation of JAZ repressor proteins by the 26S proteasome and subsequent release of activating TFs. ET binds to the receptor ETR1, and this in turn stabilizes the EIN3/EIL1 TFs which bind to the promoters of both JA and ET responsive/defense genes, activating the ERF branch of the JA defense pathway^148^.

On the other hand, to link *P. oleae* effectors with known *Phytophthora* avirulent genes, for instance, we could not find a homologue of *P. sojae* RxLR effector avh94 in the *P. oleae* genome/proteome. RxLR effector avh94 was shown to interact with soybean JAZ1/2, which is a repressor of JA signaling^83^. However, by genome sequencing and annotation, *P. oleae* was shown to possess more than 350 effectors (RxLR, crinklers, elicitins, etc.) whose function is unknown and whose identity has only been confirmed by annotation pipelines.

Here we have performed simultaneously a dual RNA-seq, with a total RNA extraction, of both host (*O. europaea*) and pathogen (*P. oleae*), with or without treatment with either *C. oleophila* or *T. atroviride*-culture filtrate. The outcome of the *P. oleae* gene expression on the host has revealed a expression pattern that was modulated by either *C. oleophila* or *T. atroviride*-culture. For instance, the cluster analysis for top DEGs of elicitins (Figure 8D) revealed that g8415.t1, g17483.t1, and g8420.t1 were upregulated upon *P. oleae* infection and downregulated by the effect of either *C. oleophila* or *T. atroviride*-culture filtrate (Figure 8D). The trend of expression for the elicitin g17483.t1 as monitored by RT-qPCR confirmed that biocontrol agents constantly over time downregulated such important elicitin (Figure 8B). Opposite trend of expression was observed for the elicitins g24094.t1, g4307.t1 and g19250.t1, which were upregulated upon treatment with either *C. oleophila* or *T. atroviride*-culture filtrate and which role is unknown. The same opposite trend was observed at 72 hpi for the INF-1 elicitins (g8407.t1 and g8411.t1), known inducer of HR (Figure 8D).

Our cluster analysis on *P. oleae* effectors (Figure 8) revealed a group of Crinkler highly differentially expressed during the infection with *P. oleae* alone (e. g. g11238.t1, g8206.t1, g23109.t1, g9419.t1, g9424.t1, g22000.t1, g10898.t1, g3434.t1 and g1422.t1) but downregulated by the treatment with either *C. oleophila* or *T. atroviride*-culture filtrate (Figure 8B). RNAseq revealed the same trend for Crinkler g723.t1 and RT-qPCR confirmed downregulation of this gene overtime as a consequence of the treatment with the BCAs (Figure 10A).

The same trend was observed for the RxLR effectors. In detail, the differentially expressed genes upregulated upon infection with *P. oleae* alone and downregulated by the treatment with either *C. oleophila* or *T. atroviride*-culture filtrate, having the highest L2FC, were g3852.t1, g16251.t1, g19132.t1, g1137.t1, g6343.t1, g567.t1, g2733.t1, g19400.t1, g24837.t1, g2412.t1, g20157.t1, and g1136.t1 respectively. For the RxLR effector g24837.t1 this trend of qPCR expression remained unchanged over time (Figure 10C).

Results obtained here strongly support the hypothesis that the molecular counteraction exerted by either *C. oleophila* or *T. atroviride-*culture filtrate applied one day before *P. oleae* inoculation, reduced significantly the pathogenicity of *P. oleae* by lowering the most effective effectors/exoenzymes by a still unclear mechanism. However, despite its action was long lasting (up to 168 hpi), it did not halt but only slowed the invasion of olive drupes by *P. oleae*. Further research will be needed to increase the effectiveness and extend further the duration of the inhibitory activity exerted by agents of biological control.

In conclusion, this study unveiled the transcriptomic events of the two- and three-actors interactions of *P. oleae* with olive fruits and each of the two evaluated biocontrol agents. Genome sequencing of *P. oleae* has strongly contributed to the study of its transcriptomics and description of the annotated pathogenesis-related genes. The olive cultivar ‘Coratina’ can be regarded as a susceptible host, despite several R-genes described here were upregulated in response to the infection (compatible host), and the two biocontrol agents only temporarily restricted the growth of the pathogen inside the drupes. However, over time (up to 168 hpi) these treatments were not able to stop *P. oleae* infection progression

## 4. Methods

### 4.1. Microorganisms and plant material

Microorganisms included in this study were: the ex-type of the yeast *C. oleophila* Montrocher strain O, ii. the ascomycete *T. atroviride* strain TS and iii. the oomycete *P. oleae* isolate VK10A. C. *oleophila* strain O, *T. atroviride* strain TS and *P. oleae* isolate VK10A were sourced from the culture collection of the Molecular Plant Pathology Laboratory of the Department of Agriculture, Food and Environment of the University of Catania (Catania, Italy) and were identified in previous studies^24,149^.

‘Coratina’ olive fruits were collected from 15-year-old olive trees from an orchard in Calabria (southern Italy), in the municipality of Mirto-Crosia, province of Cosenza (DATUM WGS 84, 39°36′54.5″ N 16°46′11.7″ E). Drupes were used to both determine the incidence of the disease following post-harvest treatments and to study the defense mechanisms triggered in the presence of the pathogen in relation to the applied treatments. Overall, 540 drupes at maturation index (MI)^7^ 4 (MI4) were collected from five trees and stored for 3 hours inside a refrigerated bag while being transported to the laboratory and stored at 8°C until used in experiments.

### 4.2. Preparation of *P. oleae-*inoculum

*Phytophthora oleae* isolate VK10A was preliminary grown on V8-Agar (V8A) at 20 ± 2°C. Then, inoculum used in following tests was represented by culture fragments (size 2 mm) cut from the growing edge of seven-day-old cultures.

### 4.3. Preparation of *T. atroviride-*culture filtrate

*Trichoderma atroviride* was grown in the dark in a 500 mL Erlenmeyer flask containing 200 mL of potato dextrose broth (PDB) at pH 7.0 under constant stirring in an orbital shaker for 30 days (28 ± 2 °C, 140 rpm)^24^. After incubation, the supernatant was collected, filtered through a 13-mm/0.22-μm nylon syringe filter and used as treatment for olive drupes in following tests.

### 4.4. Preparation of *C. oleophila* cell suspension

*C. oleophila* was grown in 500 ml shake flask cultures containing nutrient yeast broth (NYDB) at 24°C for 48 hours. Cells of *C. oleophila* were collected by centrifugation at 5000×g for 10 minutes at 4°C; then, collected cells were suspended in sterile distilled water to reach the concentration to 1×10^8^ CFU/ml (a hemacytometer was used). Obtained *C. oleophila* cell suspension was used as treatment for olive drupes in following tests.

### 4.5. Evaluating the Effectiveness of *C. oleophila* and *Trichoderma*-culture filtrate in Controlling Rots of Drupes by *P. oleae*

Olive drupes were preliminarily subjected to surface-sterilization by immersion in a 0.5 % NaClO for 30s, rinsing in sterile distilled water (sdw) and air drying for 24 hours. Once dried, olive drupes were punctured once with a sterile needle in an equatorial position and a 20 µl droplet of *C. oleophila* cell suspension, *T. atroviride*-culture filtrate or sdw (control) was pipetted onto the wound. After 24 h of incubation at 21 ± 1°C, the wound of each drupe was inoculated with a 2 mm agar plug collected from a V8A culture of *P. oleae* isolate VK10A. After inoculation, drupes were incubated at 21 ± 1°C, with 80% relative humidity and a photoperiod of 16 h light / 8 h dark. Severity of *Phytophthora* rots in olive drupes was assessed at 24, 72, and 168 hours post pathogen inoculation (hpi) and rated based on the following empirical scale: (0) no symptoms; (1) 0-25% of fruit rot around inoculation point; (2) 26-50% fruit rot; (3) 50-75 % of the fruit covered by mycelium; (4) 100 % fruit covered from mycelium. These values were used to calculate the rAUDPC according to Riolo and other authors^7,150,151^. Overall, the experimental assay included the following three treatments: i. *P. oleae*-inoculated olive drupes treated with sdw (control, IDA); ii. *P. oleae*-inoculated olive drupes treated with *T. atroviride*-culture filtrate (IDB); iii. *P. oleae*-inoculated olive drupes treated with *C. olephila* cell suspension (IDC). The experimental assay was designed as a completely randomized block, with each treatment comprising three biological replicates, each consisting of 20 drupes.

### 4.6. Assessing the experimental assay to study the Transcriptome of olive drupes and *P. oleae* in the system Olive Drupe-*P. oleae*-*C. oleophila*/*T. atroviride*-culture filtrate

Olive drupes were preliminary surface sterilized, treated with either *C. oleophila* cell suspension or *T. atroviride*-culture filtrate and *P. oleae*-inoculated as described in 4.5. Then, drupes were incubated at 21 ± 1°C, with 80% relative humidity and a photoperiod of 16 h light / 8 h dark. The experimental assay included the following seven treatments: (i) non-wounded drupes treated with 20 µl of sdw (ID1); (ii) wounded drupes treated with 20 µl of sdw (ID2); (iii) wounded drupes inoculated with *P. oleae* (ID3); (iv) wounded drupes treated with *T. atroviride-*culture filtrate and inoculated with *P. oleae* (ID4); (v) wounded drupes treated with *C. oleophila* and inoculated with *P. oleae* (ID5); (vi) wounded drupes treated with *T. atroviride*-culture filtrate (ID6); (vii) wounded drupes treated with *C. oleophila* (ID7). The experimental assay was designed as a completely randomized block, with each treatment comprising three biological replicates, each consisting of 20 drupes. In order to proceed to transcriptomic analyses, at each of the above-mentioned time points, fresh olive fragments from each treatment were collected and immediately frozen in liquid nitrogen and stored at - 80°C.

### 4.7. RNA isolation from infected olive drupes and cDNA synthesis for RT-PCR

Total RNA was extracted using RNeasy Plant Mini Kit (Qiagen, Hilden, Germany). Drupes from each treatment (100 mg) were ground to a fine powder with liquid nitrogen, following the manufacturer’s protocol and treated with TURBO DNA-free Kit (ThermoFisher Scientific, Italy). The RNA concentration was then adjusted to 200 ng/µl, and its quality was verified by performing denaturing RNA electrophoresis gel in TAE agarose^152^. First-strand cDNA was synthesized with an oligo d(T)21 primer and reverse transcription was performed using High-Capacity cDNA Reverse Transcription Kit (Applied Biosystems, Foster City, CA, United States) according to the manufacturer’s instructions.

### 4.8. Quantitative Real Time-PCR (RT-qPCR) Analysis of Gene Expression

The RT-qPCR for validating expression levels of selected genes of olive drupes and *P. oleae* (Supplementary Table 13) was performed using the QuantGene 9600 system (Bioer Technology, Hangzhou, China). Reactions were performed in a total volume of 20 μl with 10 ng of cDNA, 1 μl of 10 μM of each primer, and 10 μl of PowerUp™ SYBR™ Green Master Mix (2X, Applied Biosystems™, Foster City, CA, United States). For each treatment, three biological replicates were analysed, with triplicate RT-qPCR experiments per replicate. Thermocycling conditions included 2 min at 50°C for UDG activation, 2 min at 95°C (Dual-Lock™ DNA polymerase), followed by 40 cycles of two steps: 95°C for 15 s (denaturation) and 60°C (annealing/extension) for 1 min. Expression levels of selected genes were calculated using the 2^–ΔΔCt^ method^153^, where ΔΔCt = (Ct of target gene − Ct of reference gene)sample − (Ct of target gene − Ct of reference gene)calibrator and Ct is the threshold cycle of each transcript, defined as the point at which the amount of amplified target reaches a fixed threshold above the background fluorescence. *O. europaea* tubulin alpha-3 chain (LOC111371391, XM_022993342.1) was used as an internal reference gene. For *P. oleae*, the tubulin beta chain (g8215.t2 (TRINITY_DN6095_c0_g1_i1) gene expression has been used to normalize the RT-qPCR data. The relative expression level of each gene obtained through RT-qPCR was compared to the RNA-seq data. Expression values of the 2^−ΔΔCt^ were represented as boxplots using Python script wit "seaborn”, “pandas” and “matplotlib.pyplot” libraries. Dunnett’s test was used to calculate the significance (*p-value*) relative to the expression level (2^–ΔΔCt^) of each sample to wounded olive fruit condition by computing a Student’s t-statistic for each experimental conditions and time points (hpi).

### 4.9. Total RNA preparation and Illumina mRNA sequencing

Withdrawal of plant-fungal-oomycete material was carried out around the wounded area from the non-infected and infected drupes. The material was pulverized in liquid nitrogen and stored at -80°C until RNA extraction. Total RNA extraction was performed using the Direct-zol RNA Miniprep kit (Zymo Research, Nordic BioSite, Sweden) with a modified protocol. Frozen samples were extracted with 1.5 ml of TRI Reagent (Zymo Research, Nordic BioSite, Sweden) using ZR BashingBead Lysis (0.5 mm Y2O3 Stabilized Zirconia Beads; Zymo Research, Nordic BioSite, Sweden) and the FastPrep-24™ 5G bead beating grinder and lysis system (MP Biomedicals, Fischer Scientific, Denmark). After homogenizing, samples were spin at 21000 ×g for 1 minute and then, phenol:chloroform:isoamyl alcohol 25:24:1 (Sigma-Aldrich, Merck, Germany) extraction was performed. Further steps were performed according to the Direct-zol RNA Miniprep kit. Eluted total RNA quality check was performed using a 2100 Bioanalyzer system (Agilent, Denmark). Two biological samples were extracted in technical triplicate and samples with RNA Integrity Number (RIN) above 8 were selected for RNAseq. Samples were sequenced at Novogene (Cambridge, UK) with Illumina hiseq 4000 and 150 pair end read length (PE), at 30 million of read depths. Data quality control revealed a Q30 [(Base count of Phred value > 30) / (Total base count)] average of 93% for a total of 98% of clean reads.

### 4.10. *De novo* mRNA analysis

Illumina 150 PE, 30 M reads fastq^154^ files were processed for removal of the adapters with Trimmomatic Version 0.39^155^. For maximizing and diversifying the mRNA expression of *Phytophthora oleae*, the Trinity *de novo* mRNA assembly algorithm^156^ was applied to RNA samples of cultures of the isolate VK10A grown in three different substrates, including CZAPEK, MEA and PDA. Annotation of both trinity derived cDNA and AUGUSTUS predicted genes was performed against using Mercator4v6^157^ analysis cross-evaluation.

AUGUSTUS web prediction gave 20547 proteins of which only 12117 were containing unique scaffold represented of more than 75% as protein identity, the remaining not unique proteins represented alternate proteins or differently spliced or start (ATG) derived proteins. The same number of proteins, 20094 was found by using Trinity ORF prediction software unfiltered and after removing duplicates. Filtering the Trinity accessions for proteins capable of finding a blast hit with other Phytophthora species, then, the real number of de novo mRNA transcript was down to 13373 proteins. It needs to be considered that the Trinity prediction include partial protein/mRNA sequences due to incomplete full-length cDNA. The fasta/excel files related to AUGUSTUS and Trinity protein/CDS/mRNA prediction and annotation can be found at Zenodo (https://zenodo.org/records/11092245).

### 4.11. RNAseq data analysis

The raw Illumina sequence reads of fastaq format were processed as above by removing the adapters. The reference genome and gene model annotation files of *O. europaea* var. *sylvestris* were downloaded from NCBI (*Olea europaea* var. *sylvestris* reference genome O_europaea_v1, accession number GCF_002742605.1) and *P. oleae* reference genome VK10A, from Zenodo (https://zenodo.org/records/11092245). The filtered fastaq files were aligned to the reference genome using Hisat2 v2.0.5^158^. The reads with low alignment quality values or unpaired alignments, and alignments to multiple regions of the genome were filtered out by using samtools^159^ to generate duplicate checked and sorted binary BAM files of mapped reads to the genome. The number of reads and fragments per kilobase per million reads (FPKM) for each gene was calculated using Stringtie v2.2.0^160^. Differential gene expression analysis between the pathogen-inoculated and control samples at different time points was performed in R studio using the DESeq2 R package in a script (https://bioconductor.org/packages/DESeq2/) customized on the basis of the total normalized reads counts^161^. Corrections for false positives and false negatives were made using the Benjamini-Hochberg procedures^162^ to generate the false discovery rate (FDR).

Fold changes (L2FC, log2Ratio) were estimated from normalized gene expression levels in each sample by using the ratio of the sample Log2 of FPKM and the wounded olive fruit Log2 of FPMK as denominator. The mapped genes with an adjusted false discovery rate (FDR) ≤0.05 and |log2(FoldChange)| > 1 were considered differentially expressed, from L2FC < 0.5 were considered downregulated. To reduce the amount of differentially expressed genes according to 0.05 and 0.001 FDR, shrinkage of log2 (Fold Change) was performed using the R ’apeglm’ function^163^. Differentially expressed genes (DEGs) clustering heatmaps were generated by the pheatmap and clusterProfiler R packages^164^. Multiple correlation of the variance of L2FC for all the sample was analyzed by principal components analysis (PCA).

### 4.12. LC-MS identification of proteins in protein filtrate

*T. atroviride* strain TS was cultured *in vitro* in a YPD liquid media supplemented with malt extract to 1%. After three days, the cultures (in triplicate) were filtered through a Whatman® qualitative filter paper, Grade 1 (WHA1001325 Sigma-Aldrich, Merck, Germany) to retain the mycelia. The filtrate was further centrifuged at 20,000 ×g for 15 minutes then sterile filtered through 0.2 µm vacuum filters. The protein preparation for proteomics including digestion by trypsin and the LC-MS conditions and identification by Waters PLGS software has been described previously^165^.

## CRediT author statement

**Sebastiano Conti Taguali:** Investigation, Methodology, Formal analysis, Data Curation, Writing - Original Draft, Writing - Review & Editing

**Mario Riolo:** Conceptualization, Investigation, Methodology, Formal analysis, Software, Writing - Original Draft, Validation, Writing - Review & Editing, Visualization, Data Curation.

**Federico La Spada:** Conceptualization, Investigation, Methodology, Writing - Review & Editing.

**Giuseppe Dionisio:** Conceptualization, Validation, Data Curation, Methodology, Formal analysis, Writing - Original Draft, Writing - Review & Editing, Resources, Formal analysis, Supervision, Project administration.

**Santa Olga Cacciola**: Conceptualization, Writing - Original Draft, Validation, Writing - Review & Editing, Resources, Supervision, Funding acquisition, Project administration.

## FUNDS

This study was supported by the University of Catania, Italy, “Investigation of hytopathological problems of the main Sicilian productive contexts and eco-sustainable defense strategies (MEDIT-ECO)”-PiaCeRi-PIAno di inCEntivi per la Ricerca di Ateneo 2020-22 linea 2” “5A722192155”; the project “Smart and innovative packaging, postharvest rot management, and shipping of organic citrus fruit (BiOrangePack)” under Partnership for Research and Innovation in the Mediterranean Area (PRIMA) – H2020 (E69C20000130001); the European Union (NextGeneration EU), through the MUR-PNRR project SAMOTHRACE (ECS00000022) and Ministero dell’agricoltura, della sovranità alimentare e delle foreste (MASAF), project title: Difesa degli Agrumeti Italiani dal Malsecco – AGRIVITA (CUP: C83C23000650006).

## Data Availability Statement

Raw data for this study will be available to researchers upon reasonable request to the corresponding authors.

## Abbreviations

BCA: biological control agent
PAMP: pathogen-associated molecular pattern
ETI: effector triggered immunity
ROS: reactive oxygen species
PRs: pathogenesis related proteins.

