## Supplementary material for "Exploring the Impact of Various Treatments on Gene Expression in Olive (*Olea europaea* L.) Drupes Affected by *Phytophthora oleae*: Insights from RNA sequencing-based transcriptome analysis": Supplementary Figure 1 - PR proteins (L2FC).pdf

**Supplementary Figure 1.** ***Olea europaea*** **pathogenic related (PR) proteins cluster analysis of their L2FC.** The "wound" condition has been normalized to "not wound", while the rest of conditions has been normalized to the "wounded" condition.

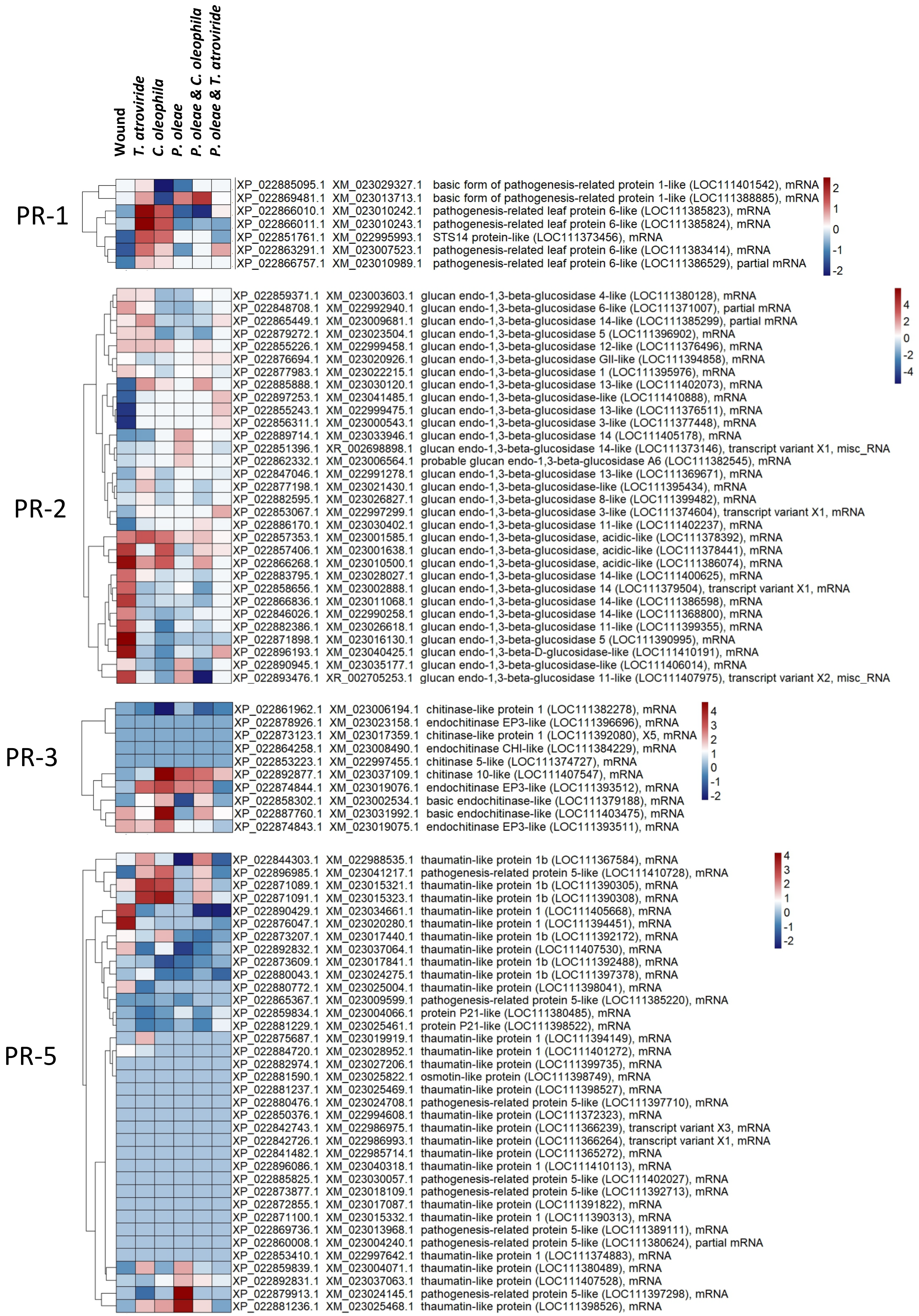

PR-6

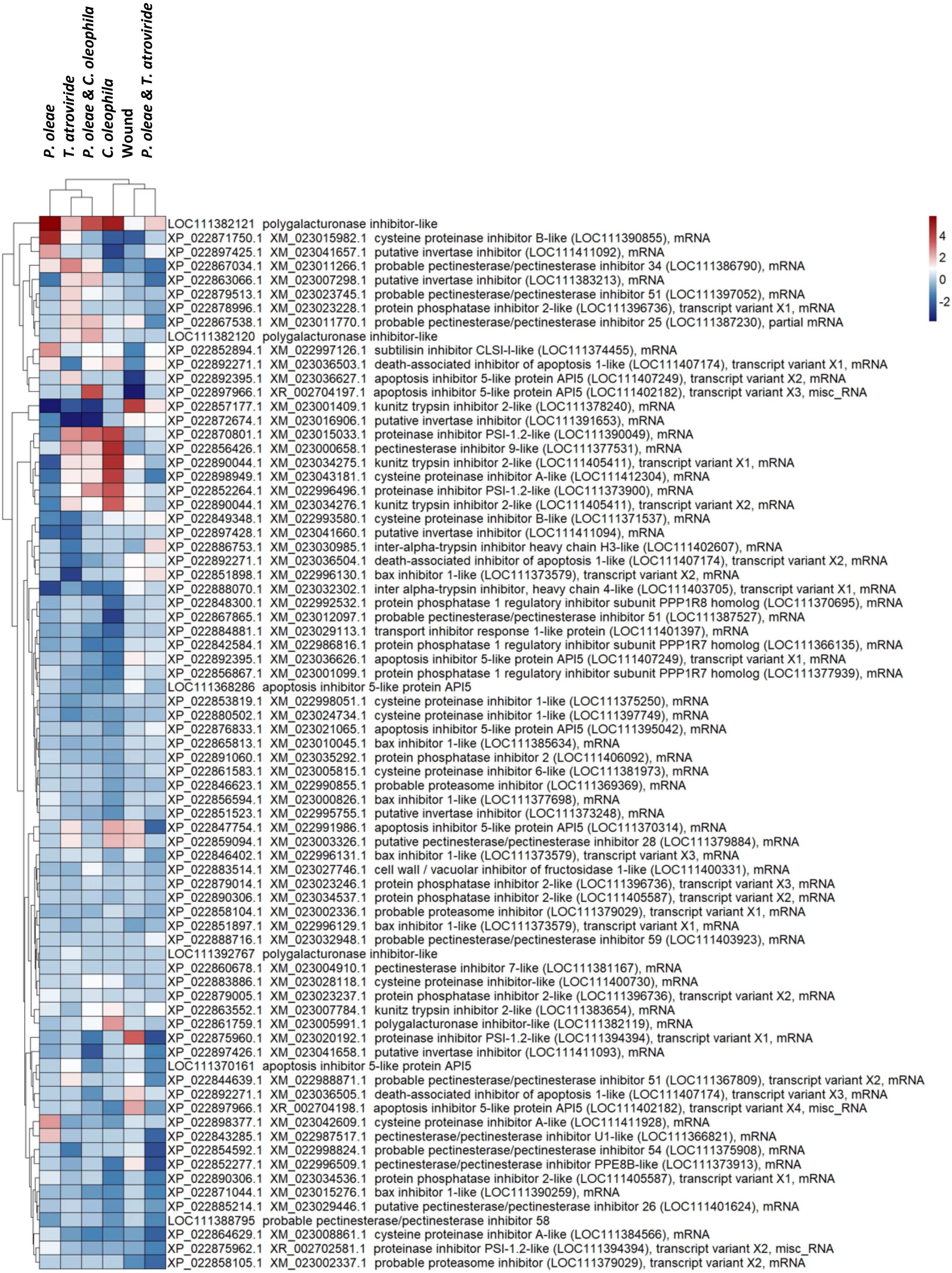
P. oleae
T. atroviride
P. oleae & C. oleophila
C. oleophila
Wound
P. oleae & T. atroviride
4
2
0
-2
LOC111382121 polygalacturonase inhibitor-like
XP_022871750.1 XM_023015982.1 cysteine proteinase inhibitor B-like (LOC111390855), mRNA
XP_022897425.1 XM_023041657.1 putative invertase inhibitor (LOC111411092), mRNA
XP_022867034.1 XM_023011266.1 probable pectinesterase/pectinesterase inhibitor 34 (LOC111386790), mRNA
XP_022863066.1 XM_023007298.1 putative invertase inhibitor (LOC111383213), mRNA
XP_022879513.1 XM_023023745.1 probable pectinesterase/pectinesterase inhibitor 51 (LOC111397052), mRNA
XP_022878996.1 XM_023023228.1 protein phosphatase inhibitor 2-like (LOC111396736), transcript variant X1, mRNA
XP_022867538.1 XM_023011770.1 probable pectinesterase/pectinesterase inhibitor 25 (LOC111387230), partial mRNA
LOC111382120 polygalacturonase inhibitor-like
XP_022852894.1 XM_022997126.1 subtilisin inhibitor CLSI-I-like (LOC111374455), mRNA
XP_022892271.1 XM_023036503.1 death-associated inhibitor of apoptosis 1-like (LOC111407174), transcript variant X1, mRNA
XP_022892395.1 XM_023036627.1 apoptosis inhibitor 5-like protein API5 (LOC111407249), transcript variant X2, mRNA
XP_022897966.1 XR_002704197.1 apoptosis inhibitor 5-like protein API5 (LOC111402182), transcript variant X3, misc_RNA
XP_022857177.1 XM_023001409.1 kunitz trypsin inhibitor 2-like (LOC111378240), mRNA
XP_022872674.1 XM_023016906.1 putative invertase inhibitor (LOC111391653), mRNA
XP_022870801.1 XM_023015033.1 proteinase inhibitor PSI-1.2-like (LOC111390049), mRNA
XP_022856426.1 XM_023000658.1 pectinesterase inhibitor 9-like (LOC111377531), mRNA
XP_022890044.1 XM_023034275.1 kunitz trypsin inhibitor 2-like (LOC111405411), transcript variant X1, mRNA
XP_022898949.1 XM_023043181.1 cysteine proteinase inhibitor A-like (LOC111412304), mRNA
XP_022852264.1 XM_022996496.1 proteinase inhibitor PSI-1.2-like (LOC111373900), mRNA
XP_022890044.1 XM_023034276.1 kunitz trypsin inhibitor 2-like (LOC111405411), transcript variant X2, mRNA
XP_022849348.1 XM_022993580.1 cysteine proteinase inhibitor B-like (LOC111371537), mRNA
XP_022897428.1 XM_023041660.1 putative invertase inhibitor (LOC111411094), mRNA
XP_022886753.1 XM_023030985.1 inter-alpha-trypsin inhibitor heavy chain H3-like (LOC111402607), mRNA
XP_022892271.1 XM_023036504.1 death-associated inhibitor of apoptosis 1-like (LOC111407174), transcript variant X2, mRNA
XP_022851898.1 XM_022996130.1 bax inhibitor 1-like (LOC111373579), transcript variant X2, mRNA
XP_022888070.1 XM_023032302.1 inter alpha-trypsin inhibitor, heavy chain 4-like (LOC111403705), transcript variant X1, mRNA
XP_022848300.1 XM_022992532.1 protein phosphatase 1 regulatory inhibitor subunit PPP1R8 homolog (LOC111370695), mRNA
XP_022867865.1 XM_023012097.1 probable pectinesterase/pectinesterase inhibitor 51 (LOC111387527), mRNA
XP_022884881.1 XM_023029113.1 transport inhibitor response 1-like protein (LOC111401397), mRNA
XP_022842584.1 XM_022986816.1 protein phosphatase 1 regulatory inhibitor subunit PPP1R7 homolog (LOC111366135), mRNA
XP_022892395.1 XM_023036626.1 apoptosis inhibitor 5-like protein API5 (LOC111407249), transcript variant X1, mRNA
XP_022856867.1 XM_023001099.1 protein phosphatase 1 regulatory inhibitor subunit PPP1R7 homolog (LOC111377939), mRNA
LOC111368286 apoptosis inhibitor 5-like protein API5
XP_022853819.1 XM_022998051.1 cysteine proteinase inhibitor 1-like (LOC111375250), mRNA
XP_022880502.1 XM_023024734.1 cysteine proteinase inhibitor 1-like (LOC111397749), mRNA
XP_022876833.1 XM_023021065.1 apoptosis inhibitor 5-like protein API5 (LOC111395042), mRNA
XP_022865813.1 XM_023010045.1 bax inhibitor 1-like (LOC111385634), mRNA
XP_022891060.1 XM_023035292.1 protein phosphatase inhibitor 2 (LOC111406092), mRNA
XP_022861583.1 XM_023005815.1 cysteine proteinase inhibitor 6-like (LOC111381973), mRNA
XP_022846623.1 XM_022990855.1 probable proteasome inhibitor (LOC111369369), mRNA
XP_022856594.1 XM_023000826.1 bax inhibitor 1-like (LOC111377698), mRNA
XP_022851523.1 XM_022995755.1 putative invertase inhibitor (LOC111373248), mRNA
XP_022847754.1 XM_022991986.1 apoptosis inhibitor 5-like protein API5 (LOC111370314), mRNA
XP_022859094.1 XM_023003326.1 putative pectinesterase/pectinesterase inhibitor 28 (LOC111379884), mRNA
XP_022846402.1 XM_022996131.1 bax inhibitor 1-like (LOC111373579), transcript variant X3, mRNA
XP_022883514.1 XM_023027746.1 cell wall / vacuolar inhibitor of fructosidase 1-like (LOC111400331), mRNA
XP_022879014.1 XM_023023246.1 protein phosphatase inhibitor 2-like (LOC111396736), transcript variant X3, mRNA
XP_022890306.1 XM_023034537.1 protein phosphatase inhibitor 2-like (LOC111405587), transcript variant X2, mRNA
XP_022858104.1 XM_023002336.1 probable proteasome inhibitor (LOC111379029), transcript variant X1, mRNA
XP_022851897.1 XM_022996129.1 bax inhibitor 1-like (LOC111373579), transcript variant X1, mRNA
XP_022888716.1 XM_023032948.1 probable pectinesterase/pectinesterase inhibitor 59 (LOC111403923), mRNA
LOC111392767 polygalacturonase inhibitor-like
XP_022860678.1 XM_023004910.1 pectinesterase inhibitor 7-like (LOC111381167), mRNA
XP_022883886.1 XM_023028118.1 cysteine proteinase inhibitor-like (LOC111400730), mRNA
XP_022879005.1 XM_023023237.1 protein phosphatase inhibitor 2-like (LOC111396736), transcript variant X2, mRNA
XP_022863552.1 XM_023007784.1 kunitz trypsin inhibitor 2-like (LOC111383654), mRNA
XP_022861759.1 XM_023005991.1 polygalacturonase inhibitor-like (LOC111382119), mRNA
XP_022875960.1 XM_023020192.1 proteinase inhibitor PSI-1.2-like (LOC111394394), transcript variant X1, mRNA
XP_022897426.1 XM_023041658.1 putative invertase inhibitor (LOC111411093), mRNA
LOC111370161 apoptosis inhibitor 5-like protein API5
XP_022844639.1 XM_022988871.1 probable pectinesterase/pectinesterase inhibitor 51 (LOC111367809), transcript variant X2, mRNA
XP_022892271.1 XM_023036505.1 death-associated inhibitor of apoptosis 1-like (LOC111407174), transcript variant X3, mRNA
XP_022897966.1 XR_002704198.1 apoptosis inhibitor 5-like protein API5 (LOC111402182), transcript variant X4, misc_RNA
XP_022898377.1 XM_023042609.1 cysteine proteinase inhibitor A-like (LOC111411928), mRNA
XP_022843285.1 XM_022987517.1 pectinesterase/pectinesterase inhibitor U1-like (LOC111366821), mRNA
XP_022854592.1 XM_022998824.1 probable pectinesterase/pectinesterase inhibitor 54 (LOC111375908), mRNA
XP_022852277.1 XM_022996509.1 pectinesterase/pectinesterase inhibitor PPE8B-like (LOC111373913), mRNA
XP_022890306.1 XM_023034536.1 protein phosphatase inhibitor 2-like (LOC111405587), transcript variant X1, mRNA
XP_022871044.1 XM_023015276.1 bax inhibitor 1-like (LOC111390259), mRNA
XP_022885214.1 XM_023029446.1 putative pectinesterase/pectinesterase inhibitor 26 (LOC111401624), mRNA
LOC111388795 probable pectinesterase/pectinesterase inhibitor 58
XP_022864629.1 XM_023008861.1 cysteine proteinase inhibitor A-like (LOC111384566), mRNA
XP_022875962.1 XR_002702581.1 proteinase inhibitor PSI-1.2-like (LOC111394394), transcript variant X2, misc_RNA
XP_022858105.1 XM_023002337.1 probable proteasome inhibitor (LOC111379029), transcript variant X2, mRNA

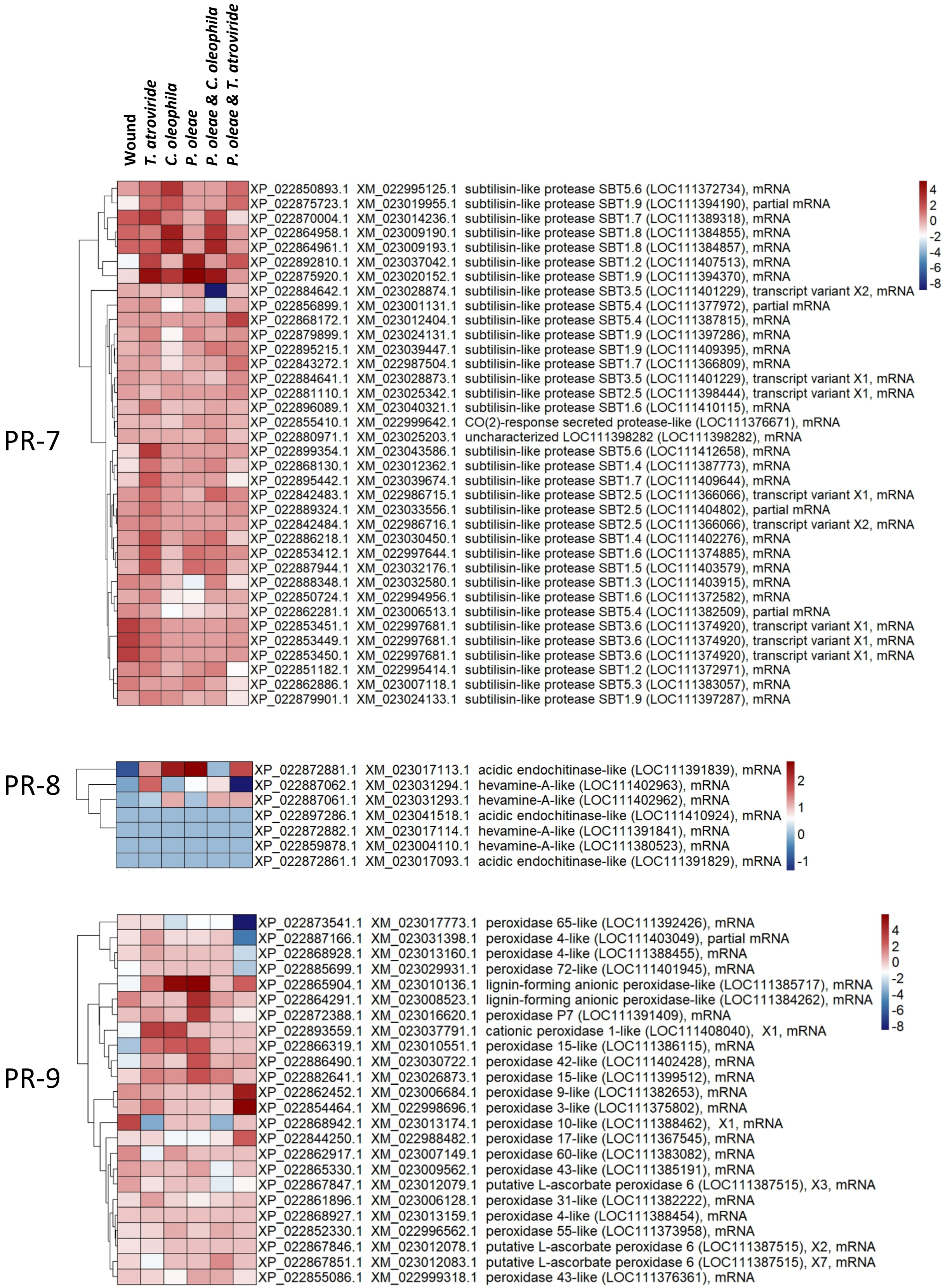
Wound
T. atroviride
C. oleophila
P. oleae
P. oleae & C. oleophila
P. oleae & T. atroviride
PR-7
XP_022850893.1 XM_022995125.1 subtilisin-like protease SBT5.6 (LOC111372734), mRNA
XP_022875723.1 XM_023019955.1 subtilisin-like protease SBT1.9 (LOC111394190), partial mRNA
XP_022870004.1 XM_023014236.1 subtilisin-like protease SBT1.7 (LOC111389318), mRNA
XP_022864958.1 XM_023009190.1 subtilisin-like protease SBT1.8 (LOC111384855), mRNA
XP_022864961.1 XM_023009193.1 subtilisin-like protease SBT1.8 (LOC111384857), mRNA
XP_022892810.1 XM_023037042.1 subtilisin-like protease SBT1.2 (LOC111407513), mRNA
XP_022875920.1 XM_023020152.1 subtilisin-like protease SBT1.9 (LOC111394370), mRNA
XP_022884642.1 XM_023028874.1 subtilisin-like protease SBT3.5 (LOC111401229), transcript variant X2, mRNA
XP_022856899.1 XM_023001131.1 subtilisin-like protease SBT5.4 (LOC111377972), partial mRNA
XP_022868172.1 XM_023012404.1 subtilisin-like protease SBT5.4 (LOC111387815), mRNA
XP_022879899.1 XM_023024131.1 subtilisin-like protease SBT1.9 (LOC111397286), mRNA
XP_022895215.1 XM_023039447.1 subtilisin-like protease SBT1.9 (LOC111409395), mRNA
XP_022843272.1 XM_022987504.1 subtilisin-like protease SBT1.7 (LOC111366809), mRNA
XP_022884641.1 XM_023028873.1 subtilisin-like protease SBT3.5 (LOC111401229), transcript variant X1, mRNA
XP_022881110.1 XM_023025342.1 subtilisin-like protease SBT2.5 (LOC111398444), transcript variant X1, mRNA
XP_022896089.1 XM_023040321.1 subtilisin-like protease SBT1.6 (LOC111410115), mRNA
XP_022855410.1 XM_022999642.1 CO(2)-response secreted protease-like (LOC111376671), mRNA
XP_022880971.1 XM_023025203.1 uncharacterized LOC111398282 (LOC111398282), mRNA
XP_022899354.1 XM_023043586.1 subtilisin-like protease SBT5.6 (LOC111412658), mRNA
XP_022868130.1 XM_023012362.1 subtilisin-like protease SBT1.4 (LOC111387773), mRNA
XP_022895442.1 XM_023039674.1 subtilisin-like protease SBT1.7 (LOC111409644), mRNA
XP_022842483.1 XM_022986715.1 subtilisin-like protease SBT2.5 (LOC111366066), transcript variant X1, mRNA
XP_022889324.1 XM_023033556.1 subtilisin-like protease SBT2.5 (LOC111404802), partial mRNA
XP_022842484.1 XM_022986716.1 subtilisin-like protease SBT2.5 (LOC111366066), transcript variant X2, mRNA
XP_022886218.1 XM_023030450.1 subtilisin-like protease SBT1.4 (LOC111402276), mRNA
XP_022853412.1 XM_022997644.1 subtilisin-like protease SBT1.6 (LOC111374885), mRNA
XP_022887944.1 XM_023032176.1 subtilisin-like protease SBT1.5 (LOC111403579), mRNA
XP_022888348.1 XM_023032580.1 subtilisin-like protease SBT1.3 (LOC111403915), mRNA
XP_022850724.1 XM_022994956.1 subtilisin-like protease SBT1.6 (LOC111372582), mRNA
XP_022862281.1 XM_023006513.1 subtilisin-like protease SBT5.4 (LOC111382509), partial mRNA
XP_022853451.1 XM_022997681.1 subtilisin-like protease SBT3.6 (LOC111374920), transcript variant X1, mRNA
XP_022853449.1 XM_022997681.1 subtilisin-like protease SBT3.6 (LOC111374920), transcript variant X1, mRNA
XP_022853450.1 XM_022997681.1 subtilisin-like protease SBT3.6 (LOC111374920), transcript variant X1, mRNA
XP_022851182.1 XM_022995414.1 subtilisin-like protease SBT1.2 (LOC111372971), mRNA
XP_022862886.1 XM_023007118.1 subtilisin-like protease SBT5.3 (LOC111383057), mRNA
XP_022879901.1 XM_023024133.1 subtilisin-like protease SBT1.9 (LOC111397287), mRNA
4
2
0
-2
-4
-6
-8
PR-8
XP_022872881.1 XM_023017113.1 acidic endochitinase-like (LOC111391839), mRNA
XP_022887062.1 XM_023031294.1 hevamine-A-like (LOC111402963), mRNA
XP_022887061.1 XM_023031293.1 hevamine-A-like (LOC111402962), mRNA
XP_022897286.1 XM_023041518.1 acidic endochitinase-like (LOC111410924), mRNA
XP_022872882.1 XM_023017114.1 hevamine-A-like (LOC111391841), mRNA
XP_022859878.1 XM_023004110.1 hevamine-A-like (LOC111380523), mRNA
XP_022872861.1 XM_023017093.1 acidic endochitinase-like (LOC111391829), mRNA
2
1
0
-1
PR-9
XP_022873541.1 XM_023017773.1 peroxidase 65-like (LOC111392426), mRNA
XP_022887166.1 XM_023031398.1 peroxidase 4-like (LOC111403049), partial mRNA
XP_022868928.1 XM_023013160.1 peroxidase 4-like (LOC111388455), mRNA
XP_022885699.1 XM_023029931.1 peroxidase 72-like (LOC111401945), mRNA
XP_022865904.1 XM_023010136.1 lignin-forming anionic peroxidase-like (LOC111385717), mRNA
XP_022864291.1 XM_023008523.1 lignin-forming anionic peroxidase-like (LOC111384262), mRNA
XP_022872388.1 XM_023016620.1 peroxidase P7 (LOC111391409), mRNA
XP_022893559.1 XM_023037791.1 cationic peroxidase 1-like (LOC111408040), X1, mRNA
XP_022866319.1 XM_023010551.1 peroxidase 15-like (LOC111386115), mRNA
XP_022886490.1 XM_023030722.1 peroxidase 42-like (LOC111402428), mRNA
XP_022882641.1 XM_023026873.1 peroxidase 15-like (LOC111399512), mRNA
XP_022862452.1 XM_023006684.1 peroxidase 9-like (LOC111382653), mRNA
XP_022854464.1 XM_022998696.1 peroxidase 3-like (LOC111375802), mRNA
XP_022868942.1 XM_023013174.1 peroxidase 10-like (LOC111388462), X1, mRNA
XP_022844250.1 XM_022988482.1 peroxidase 17-like (LOC111367545), mRNA
XP_022862917.1 XM_023007149.1 peroxidase 60-like (LOC111383082), mRNA
XP_022865330.1 XM_023009562.1 peroxidase 43-like (LOC111385191), mRNA
XP_022867847.1 XM_023012079.1 putative L-ascorbate peroxidase 6 (LOC111387515), X3, mRNA
XP_022861896.1 XM_023006128.1 peroxidase 31-like (LOC111382222), mRNA
XP_022868927.1 XM_023013159.1 peroxidase 4-like (LOC111388454), mRNA
XP_022852330.1 XM_022996562.1 peroxidase 55-like (LOC111373958), mRNA
XP_022867846.1 XM_023012078.1 putative L-ascorbate peroxidase 6 (LOC111387515), X2, mRNA
XP_022867851.1 XM_023012083.1 putative L-ascorbate peroxidase 6 (LOC111387515), X7, mRNA
XP_022855086.1 XM_022999318.1 peroxidase 43-like (LOC111376361), mRNA
4
2
0
-2
-4
-6
-8

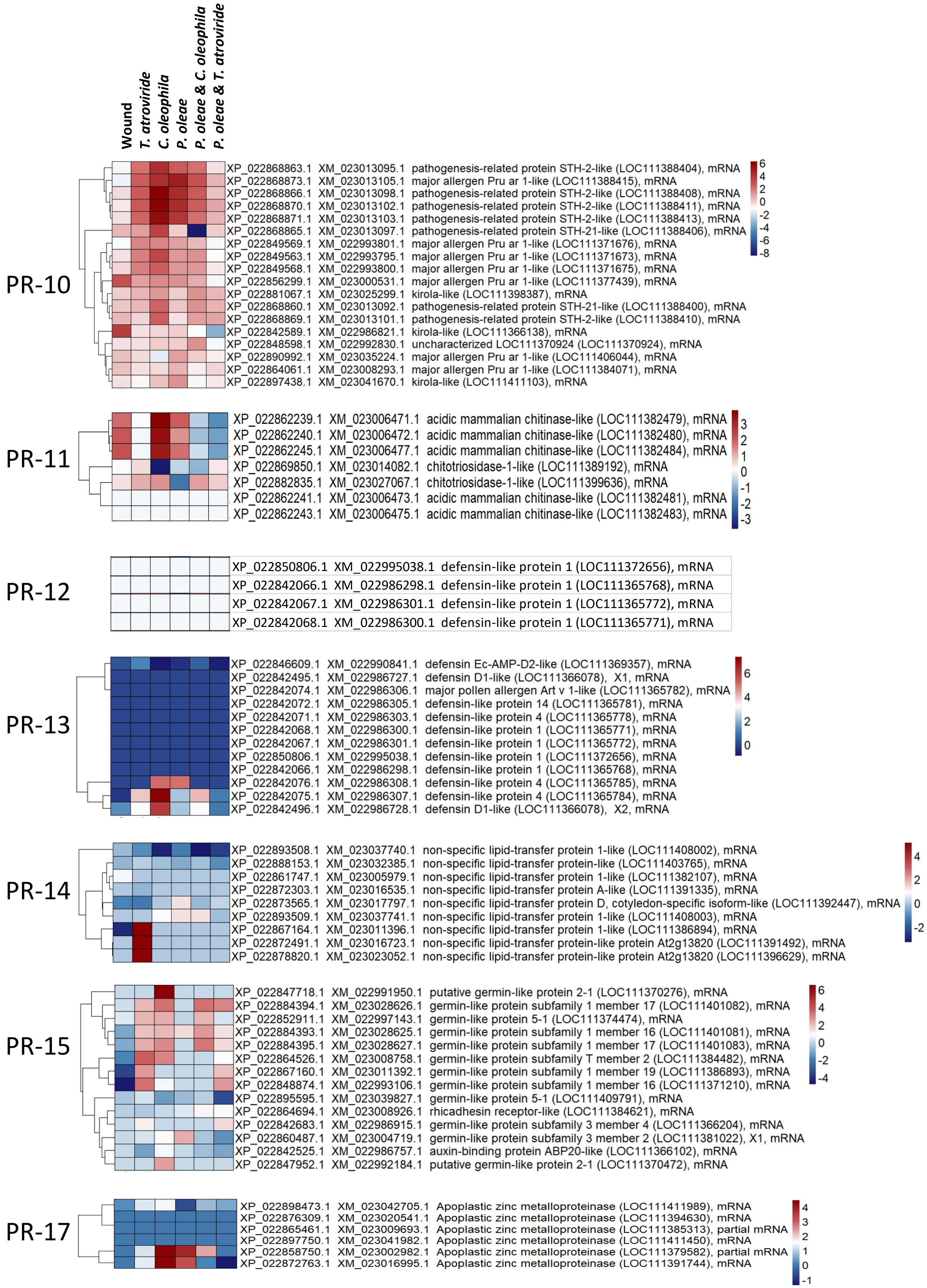

Wound
T. atroviride
C. oleophila
P. oleae
P. oleae & C. oleophila
P. oleae & T. atroviride
PR-10
XP_022868863.1 XM_023013095.1 pathogenesis-related protein STH-2-like (LOC111388404), mRNA
XP_022868873.1 XM_023013105.1 major allergen Pru ar 1-like (LOC111388415), mRNA
XP_022868866.1 XM_023013098.1 pathogenesis-related protein STH-2-like (LOC111388408), mRNA
XP_022868870.1 XM_023013102.1 pathogenesis-related protein STH-2-like (LOC111388411), mRNA
XP_022868871.1 XM_023013103.1 pathogenesis-related protein STH-2-like (LOC111388413), mRNA
XP_022868865.1 XM_023013097.1 pathogenesis-related protein STH-21-like (LOC111388406), mRNA
XP_022849569.1 XM_022993801.1 major allergen Pru ar 1-like (LOC111371676), mRNA
XP_022849563.1 XM_022993795.1 major allergen Pru ar 1-like (LOC111371673), mRNA
XP_022849568.1 XM_022993800.1 major allergen Pru ar 1-like (LOC111371675), mRNA
XP_022856299.1 XM_023000531.1 major allergen Pru ar 1-like (LOC111377439), mRNA
XP_022881067.1 XM_023025299.1 kirola-like (LOC111398387), mRNA
XP_022868860.1 XM_023013092.1 pathogenesis-related protein STH-21-like (LOC111388400), mRNA
XP_022868869.1 XM_023013101.1 pathogenesis-related protein STH-2-like (LOC111388410), mRNA
XP_022842589.1 XM_022986821.1 kirola-like (LOC111366138), mRNA
XP_022848598.1 XM_022992830.1 uncharacterized LOC111370924 (LOC111370924), mRNA
XP_022890992.1 XM_023035224.1 major allergen Pru ar 1-like (LOC111406044), mRNA
XP_022864061.1 XM_023008293.1 major allergen Pru ar 1-like (LOC111384071), mRNA
XP_022897438.1 XM_023041670.1 kirola-like (LOC111411103), mRNA
6
4
2
0
-2
-4
-6
-8
PR-11
XP_022862239.1 XM_023006471.1 acidic mammalian chitinase-like (LOC111382479), mRNA
XP_022862240.1 XM_023006472.1 acidic mammalian chitinase-like (LOC111382480), mRNA
XP_022862245.1 XM_023006477.1 acidic mammalian chitinase-like (LOC111382484), mRNA
XP_022869850.1 XM_023014082.1 chitotriosidase-1-like (LOC111389192), mRNA
XP_022882835.1 XM_023027067.1 chitotriosidase-1-like (LOC111399636), mRNA
XP_022862241.1 XM_023006473.1 acidic mammalian chitinase-like (LOC111382481), mRNA
XP_022862243.1 XM_023006475.1 acidic mammalian chitinase-like (LOC111382483), mRNA
3
2
1
0
-1
-2
-3
PR-12
XP_022850806.1 XM_022995038.1 defensin-like protein 1 (LOC111372656), mRNA
XP_022842066.1 XM_022986298.1 defensin-like protein 1 (LOC111365768), mRNA
XP_022842067.1 XM_022986301.1 defensin-like protein 1 (LOC111365772), mRNA
XP_022842068.1 XM_022986300.1 defensin-like protein 1 (LOC111365771), mRNA
PR-13
XP_022846609.1 XM_022990841.1 defensin Ec-AMP-D2-like (LOC111369357), mRNA
XP_022842495.1 XM_022986727.1 defensin D1-like (LOC111366078), X1, mRNA
XP_022842074.1 XM_022986306.1 major pollen allergen Art v 1-like (LOC111365782), mRNA
XP_022842072.1 XM_022986305.1 defensin-like protein 14 (LOC111365781), mRNA
XP_022842071.1 XM_022986303.1 defensin-like protein 4 (LOC111365778), mRNA
XP_022842068.1 XM_022986300.1 defensin-like protein 1 (LOC111365771), mRNA
XP_022842067.1 XM_022986301.1 defensin-like protein 1 (LOC111365772), mRNA
XP_022850806.1 XM_022995038.1 defensin-like protein 1 (LOC111372656), mRNA
XP_022842066.1 XM_022986298.1 defensin-like protein 1 (LOC111365768), mRNA
XP_022842076.1 XM_022986308.1 defensin-like protein 4 (LOC111365785), mRNA
XP_022842075.1 XM_022986307.1 defensin-like protein 4 (LOC111365784), mRNA
XP_022842496.1 XM_022986728.1 defensin D1-like (LOC111366078), X2, mRNA
6
4
2
0
PR-14
XP_022893508.1 XM_023037740.1 non-specific lipid-transfer protein 1-like (LOC111408002), mRNA
XP_022888153.1 XM_023032385.1 non-specific lipid-transfer protein-like (LOC111403765), mRNA
XP_022861747.1 XM_023005979.1 non-specific lipid-transfer protein 1-like (LOC111382107), mRNA
XP_022872303.1 XM_023016535.1 non-specific lipid-transfer protein A-like (LOC111391335), mRNA
XP_022873565.1 XM_023017797.1 non-specific lipid-transfer protein D, cotyledon-specific isoform-like (LOC111392447), mRNA
XP_022893509.1 XM_023037741.1 non-specific lipid-transfer protein 1-like (LOC111408003), mRNA
XP_022867164.1 XM_023011396.1 non-specific lipid-transfer protein 1-like (LOC111386894), mRNA
XP_022872491.1 XM_023016723.1 non-specific lipid-transfer protein-like protein At2g13820 (LOC111391492), mRNA
XP_022878820.1 XM_023023052.1 non-specific lipid-transfer protein-like protein At2g13820 (LOC111396629), mRNA
4
2
0
-2
PR-15
XP_022847718.1 XM_022991950.1 putative germin-like protein 2-1 (LOC111370276), mRNA
XP_022884394.1 XM_023028626.1 germin-like protein subfamily 1 member 17 (LOC111401082), mRNA
XP_022852911.1 XM_022997143.1 germin-like protein 5-1 (LOC111374474), mRNA
XP_022884393.1 XM_023028625.1 germin-like protein subfamily 1 member 16 (LOC111401081), mRNA
XP_022884395.1 XM_023028627.1 germin-like protein subfamily 1 member 17 (LOC111401083), mRNA
XP_022864526.1 XM_023008758.1 germin-like protein subfamily T member 2 (LOC111384482), mRNA
XP_022867160.1 XM_023011392.1 germin-like protein subfamily 1 member 19 (LOC111386893), mRNA
XP_022848874.1 XM_022993106.1 germin-like protein subfamily 1 member 16 (LOC111371210), mRNA
XP_022895595.1 XM_023039827.1 germin-like protein 5-1 (LOC111409791), mRNA
XP_022864694.1 XM_023008926.1 rhicadhesin receptor-like (LOC111384621), mRNA
XP_022842683.1 XM_022986915.1 germin-like protein subfamily 3 member 4 (LOC111366204), mRNA
XP_022860487.1 XM_023004719.1 germin-like protein subfamily 3 member 2 (LOC111381022), X1, mRNA
XP_022842525.1 XM_022986757.1 auxin-binding protein ABP20-like (LOC111366102), mRNA
XP_022847952.1 XM_022992184.1 putative germin-like protein 2-1 (LOC111370472), mRNA
6
4
2
0
-2
-4
PR-17
XP_022898473.1 XM_023042705.1 Apoplastic zinc metalloproteinase (LOC111411989), mRNA
XP_022876309.1 XM_023020541.1 Apoplastic zinc metalloproteinase (LOC111394630), mRNA
XP_022865461.1 XM_023009693.1 Apoplastic zinc metalloproteinase (LOC111385313), partial mRNA
XP_022897750.1 XM_023041982.1 Apoplastic zinc metalloproteinase (LOC111411450), mRNA
XP_022858750.1 XM_023002982.1 Apoplastic zinc metalloproteinase (LOC111379582), partial mRNA
XP_022872763.1 XM_023016995.1 Apoplastic zinc metalloproteinase (LOC111391744), mRNA
4
3
2
1
0
-1
