## Supplementary material for "Exploring the Impact of Various Treatments on Gene Expression in Olive (*Olea europaea* L.) Drupes Affected by *Phytophthora oleae*: Insights from RNA sequencing-based transcriptome analysis": Supplementary Figure 2 - RALF peptides homologues from fungi (final).pdf

**Supplementary Figure 2.** Proteins alignment of RALF-like peptide homologues for selected fungi (Panel A). *P. oleae* RALF-like homologues top blast hits were aligned to Fusarium and Colletotrichum RALF-like proteins. However, only the protein g3545.t1 possessed the double cysteine pattern, Kexin/Subtilisin cleavage site and FER binding motif more resembling the one present in Colletotrichum rather than the one in Fusarium (Panel B).

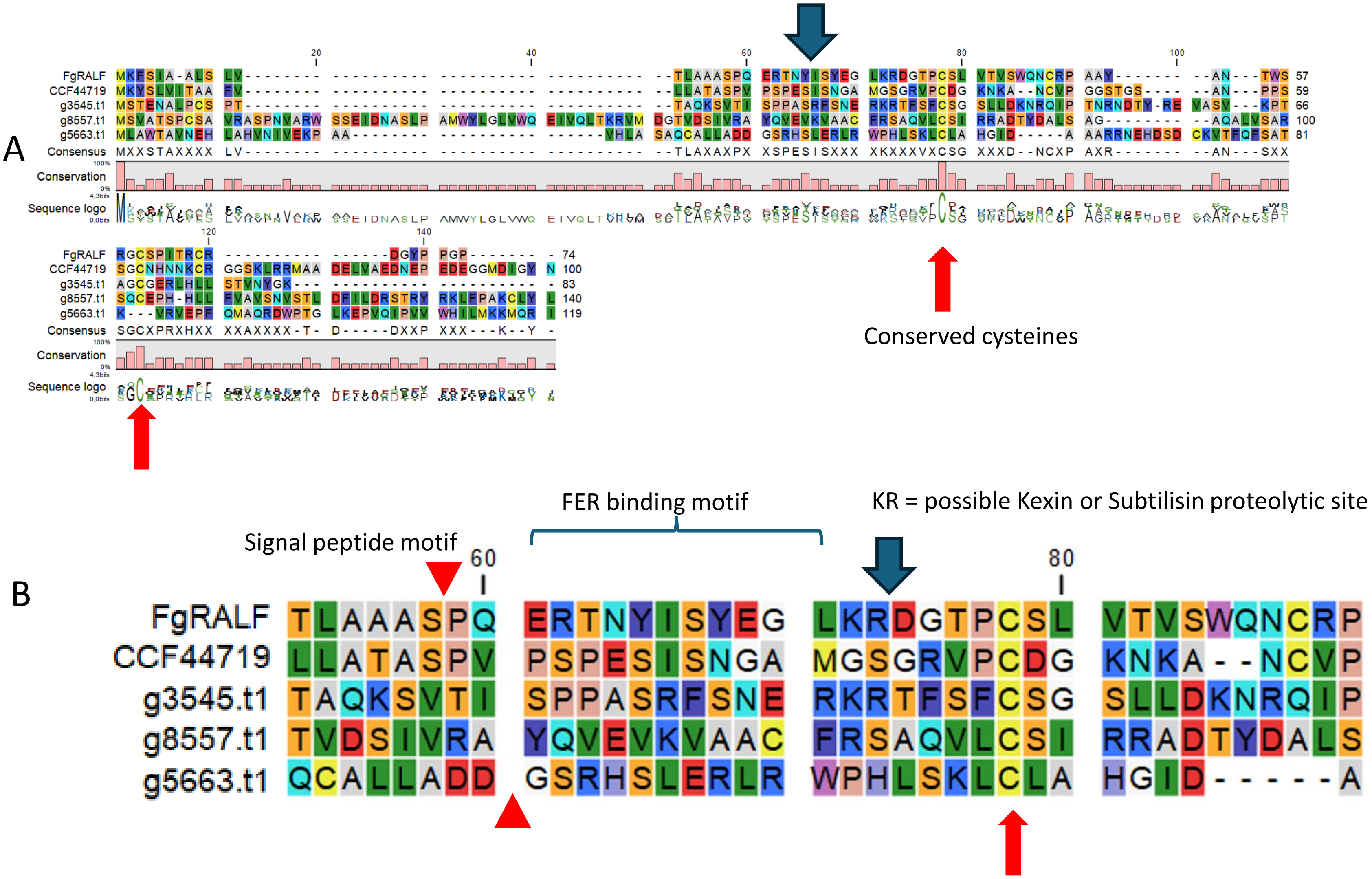


Conserved cysteines

*FgRALF= Fusarium graminearum RALF-like protein;*
*CCF44719= Colletotrichum higginsianum RALF-like protein.*
